# A giant ankyrin-B mechanism for neuro-diversity/divergence through stochastic ectopic axon projections

**DOI:** 10.1101/479741

**Authors:** Rui Yang, Kathryn K. Walder-Christensen, Namsoo Kim, Danwei Wu, Damaris Lorenzo, Alexandra Badea, Shen Gu, Haley Streff, Claudia Soler-Alfonso, Linyan Meng, William C Wetsel, Yong-Hui Jiang, Henry Yin, Vann Bennett

## Abstract

*ANK2* is a high-confidence autism spectrum disorder (ASD) gene where affected individuals exhibit diverse symptoms with a wide range of IQ. We report a cellular mechanism resulting in stochastic brain connectivity that provides a rationale for both gain and loss of function due to *ANK2* mutation. Deficiency of giant ankyrin-B (ankB), the neurospecific *ANK2* mutation target, results in ectopic CNS axon tracts associated with increased axonal branching. We elucidate a mechanism limiting axon branching, whereby giant ankB is recruited by L1CAM to periodic axonal plasma membrane domains where it coordinates cortical microtubules and prevents microtubule stabilization of nascent axon branches. Heterozygous giant ankB mutant mice exhibit innate social deficits combined with normal/enhanced cognitive function. Thus, giant ankB-deficiency results in gain of aberrant structural connectivity with penetrant behavioral consequences that may contribute to both high and low-function ASD and other forms of neurodiversity/divergence.

---

Autism spectrum disorders (ASD) are heterogeneous neurodevelopmental conditions with impaired reciprocal social behavior and restricted or repetitive behavior. ASD can coexist with intellectual disability as well as neurological co-morbidities such as seizures. However, high-functioning ASD occurs with normal or above normal IQ and may be viewed as a manifestation of neurodiversity rather than a disorder^1^. Genetic risk for ASD is conferred by variants of up to 1000 different genes, with functions that are believed, based on unbiased pathway analyses, to converge on synaptic activity and gene regulation^2^. However, global defects in synaptic function or gene regulation are unlikely to explain high-functioning ASD. Indeed, rare examples of high-functioning ASD are associated with a synapse- and brain region-specific gain-of-function due to neuroligin-3 mutation^3–5^. Other than synapses, cellular disease substrates for high-functioning ASD are not known.

Highly penetrant genes recently identified in genome-wide studies^6,7^ promise to provide new insights into the cellular origins of ASD through direct experimentation without reliance on current gene annotations. *ANK2* is unusual among these high-risk ASD genes in that mutations resulting in an ASD diagnosis occur with intellectual performance ranging from severely impaired to normal (Extended Data Table 1, Fig. 1f-g)^6,7^.

**Fig. 1.**
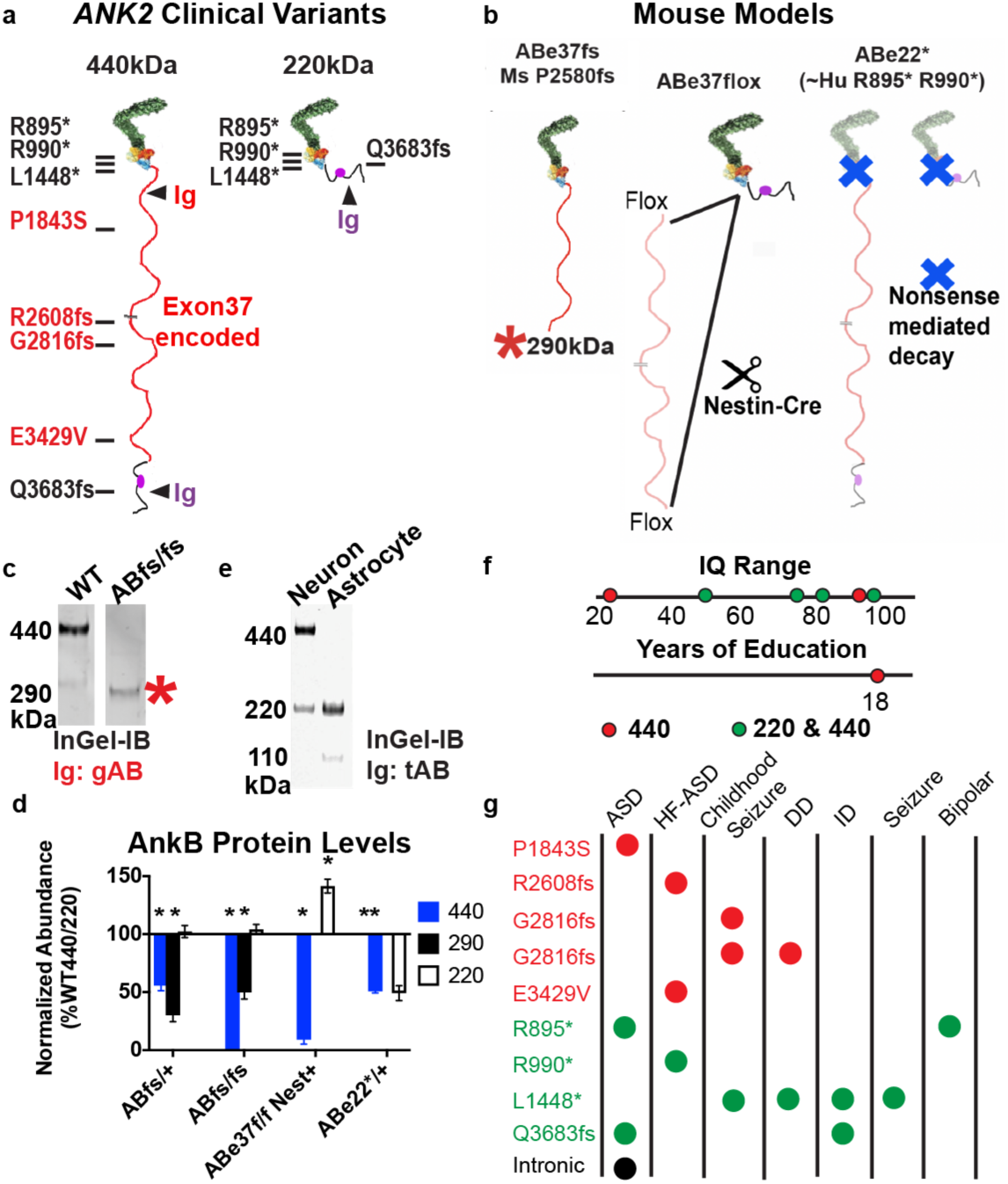
Patients with variants affecting giant ankB exhibit neurodiverse and neurodivergent phenotypes. a. Schematic of giant ankB and 220kDa ankB with *de novo* ASD variants mapped. Location of the peptides used in immunodetection are labeled with arrows. b. Scheme of resulting protein changes in mouse models of ANK2 deficiency/mutation. c. Representative in-gel immunoblot using Ig against the exon 37 encoded region. * indicates a truncated 290kDa ankB polypeptide. d. Quantification of ankB polypeptides in mouse models of ANK2 deficiency/mutation. n=3mice/genotype. e. Representative in-gel immunoblot using Ig against c-terminus of ankB in cultured neurons and astrocytes. f. IQ range and years of education for patients with *ANK2* variants. g. Diagnoses of individuals with *ANK2* variants. (f, g: red indicates mutation affects giant ankB only, green affects both 220 and giant ankB). (Mean ±s.e.m. *P<0.001, two-way ANOVA, post-hoc t-test).

*ANK2* is a member of the ankyrin gene family that, together with their spectrin partners, is responsible for functional organization of vertebrate plasma membranes in a variety of physiological contexts^8^. *ANK2* encodes 2 major ankyrin-B (ankB) polypeptides: one of 220 kDa that is broadly expressed and associated with cardiac arrhythmia^9,10^ as well as age-dependent obesity^11,12^, and another of 440 kDa (giant ankB) that is expressed only in neurons and targets to axons (Extended Data Fig.1a)^13–17^. 220-kDa ankB functions in polarized transport of intracellular organelles as a PI3P adaptor that also binds to dynactin^17,18^. Mice lacking both ankyrin polypeptides exhibit a major reduction in long axon tracts including within the corpus callosum^17,19^.

Neuro-specific giant ankB includes 2133 additional residues, likely configured as an extended polypeptide, that are not present in 220 kDa ankB, and are encoded by a single 6.4 kb exon (exon 37) acquired early in vertebrate evolution (Fig. 1a)^8^. Giant ankB is the predominant ankB isoform in early postnatal CNS development, where it partners with the membrane-spanning cell adhesion molecule L1CAM in long axon tracts^19^. Three human *ANK2* ASD mutations (R2608 frameshift and P1843S) are located in exon 37, and these mutations only affect giant ankB, while others (R895*, R990*) affect both giant ankB and 220 kDa ankB. R895* and R990* mutations likely result in haploinsufficiency based on a mouse model bearing nonsense mutation in exon 22 of ANK2 (Fig. 1b,d). IQ scores ranging from 20 to 94 do not segregate by isoform, as both the highest and lowest scores were achieved by individuals with giant ankB mutations (Fig 1f). This distribution of *ANK2* mutations and clinical history of probands bearing these mutations suggests that deficiency/mutation of giant ankB alone is sufficient to cause ASD with a wide range of intellectual ability.

We identified new *ANK2* variants without an explicit ASD diagnosis in patients with developmental delay and/or childhood seizures (Case reports, Extended data table 1, Fig. 1f-g). Two identified probands with ANK2 variants, one affecting both 220 and giant ankB (L1448*) and another affecting only the giant ankB (R2816fs), exhibit early-onset neurological symptoms including developmental delay and seizures (Fig.1f-g). The adult R2816fs individual had childhood seizures that subsequently resolved, and developmental delay, but graduated from college, and is employed (Case reports; Fig. 1f). The close proximity of the two frameshift mutations in giant ankB that produce distinct neurological phenotypes, suggests that giant ankB is not only an ASD risk gene but should more generally be classified as a neurodiversity/divergence gene.

## Abnormal CNS structural connectivity in ABe37fs/fs mice

In order to understand the consequences of giant ankB ASD mutation *in vivo*, we generated a C57Blk6 mouse strain (ABe37fs) using CRISPR to introduce a P2580) frameshift mutation in order to mimic two human *ANK2* frameshift mutations in exon 37 of *ANK2* (R2608fs; G2816fs)(Figure 1; extended table 1)^7^. ABe37fs/fs mice lack 440 kDa ankB, but instead express reduced levels of a truncated 290kDa polypeptide without effecting the levels of 220kDa ankB (Fig. 1c, Extended Data Fig. 1c). ABe37fs/fs and ABe37fs/+ mice are born at normal Mendelian ratios, are fertile, with normal body weights and lifespans (Extended Data Fig.1d).

Giant ankB collaborates with L1CAM in patterning of pre-myelinated axons during postnatal development^14,16^. In addition, Y1229H mutation in the L1CAM ankyrin-binding motif eliminates its ankyrin-binding activity and results in axon-targeting defects in mice as well as a neurodevelopmental syndrome in humans^14,20,21^. These considerations suggested that abnormal axon pathfinding may underlie the altered neurological symptoms in the *ANK2* patients. We therefore visualized overall brain morphology and axon tracts in brains of PND28 control and ABe37fs/fs mice using magnetic resonance imaging (MRI) and diffusion tensor imaging (DTI) ^22,23^. We first determined whether ABe37fs/fs mice exhibited overall structural differences in brain morphology using the same MRI data set as for DTI, since some mouse models for autism exhibit macrocephaly^2^. However, WT and ABe37fs/fs mice exhibited no statistically significant differences (false discovery rate less than 5%) in regional brain volumes after multiple comparison correction, (Extended Data Table 3).

High-functioning ASD patients exhibit increased variability in brain connectivity as well as increased interhemispheric asymmetry^24^. We therefore evaluated interhemispheric track asymmetry for either the entire brain or for the cerebral cortex by performing voxel-based comparisons within hemispheres of individual mice using tracts based on fractional anisotropy (FA) (Methods). This analysis revealed a small difference in asymmetry for the whole brain which was not statistically significant, but was consistent with overall normal brain morphology (Supplemental Table 3). However, ABe37fs/fs cortexes exhibited a significant increase in asymmetry long tracks (Fig. 2a). To visualize the tractography underlying the asymmetry, we displayed tracks passing through the cingulate area 24a, which is a cortical region located close to the midline of the brain where fibers project symmetrically to both hemispheres in WT brains. ABe37fs/fs mice exhibited increased ectopic axon tracts counted based on loss of symmetry (Fig. 2b, Extended Data Fig. 2). The projection of these ectopic tracks varied between individuals (Extended Data Fig. 2).

**Fig. 2.**
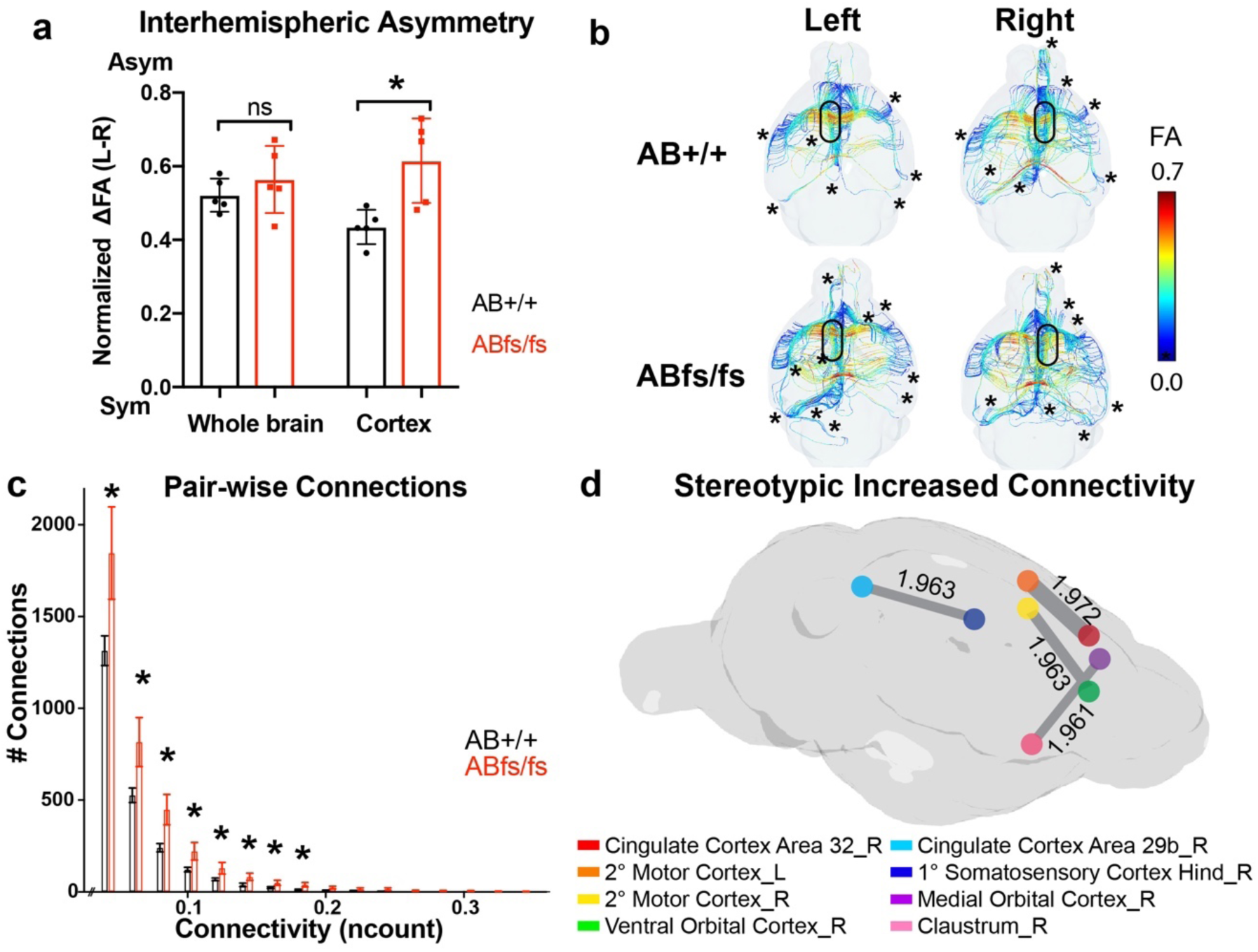
ABe37fs mice have a gain of random connectivity in the brain by DTI. a. Quantification of interhemispheric voxel asymmetry analysis (n=5 mice/genotype). b. Tracts passing through the cingulate cortex area 24a (location marked by black oval) in the left or right hemisphere. Color is based on FA value. * marks asymmetric tract. c. Histogram pair-wise connectivity from WT and ABe37fs/fs brains. d. Connectivity rendering of highest effected pairs in ABe37fs/fs brain. Absolute values of the scaled differences in connectivity are labeled. (N=5 mice/genotype; mean ±s.e.m, a. t-test. c. Kolmogorov-Smirnov test, t-test. *P<0.05).

Next, we examined whole brain pair-wise connectivity based on an atlas of 332 brain regions. Strikingly, this analysis revealed an increase in 68% of pair-wise brain connections and greater variability among animals in ABe37fs/fs mice (Fig. 2c). We next identified locations with the highest magnitudes of increased connectivity (at least 100 connecting fibers). These sites were all in the cortex where the asymmetry also is most increased (Fig. 2d). The top 4 pairs of connections that met these criteria were cingulate cortex area 32 to secondary motor cortex, medial orbital cortex to claustrum, secondary motor cortex to ventral orbital cortex, and cingulate cortex 29 to hind region of the primary somatosensory cortex (Fig. 2d). In summary, ABe37fs/fs mice exhibit overall normal brain morphology but with alteration in the cortex that include increased individual variability in structural connectivity, increased inter-hemispheric asymmetry, as well as overall increased stereotypical connectivity that is most pronounced in a subset of cortical regions.

## Giant ankB mutation causes exuberant axon branching

To resolve the cellular basis for ectopic axon tracts revealed by DTI in ABe37fs/fs brains, we next examined cultured hippocampal neurons from both ABe37fs/fs and ABe37fs/+ mice. ABe37fs/fs and ABe37fs/+ neurons consistently exhibited increased axon branching with no change in primary axon length or dendritic morphology (Fig. 3a, Extended Data Fig. 3b). This increase in axon branching in cultured neurons was restored to normal levels when ABe37fs/fs cells were transfected with giant ankB cDNA (Fig. 3a). We also observed increased axon branching in neurons from ankB exon37 floxed/Nestin-Cre mice, where giant ankB is missing (Extended Data Fig. 3a). Thus increased axonal branching in ABe37fs/fs and ABe37fs/+ neurons is a gain of cellular activity that results from loss of giant ankB.

**Fig. 3.**
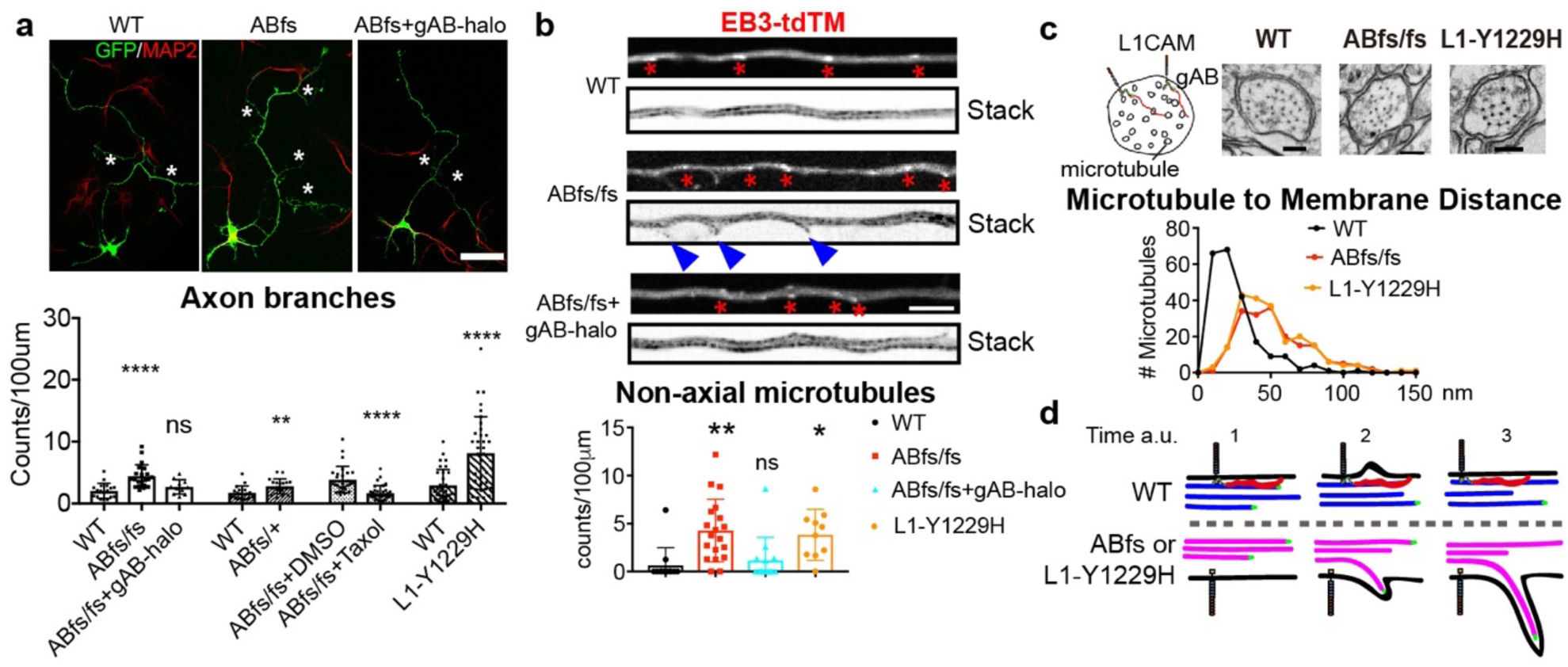
Increased axonal branching in ABe37fs/fs and L1CAM Y1229H neurons. a. GFP-transfected cultured neurons (green=cell fill, red=MAP2, axon branches highlighted by asterisks); and quantification of axon branching from the indicated conditions (n = 18, 21, 11, 18, 19, 17, 32, 32, 35 from left to right in 3 cultures, Scale bar: 20 µm). b. Axons from single-frame and time-lapse stacked images of EB3-tdTM transfected neurons (n= 12, 19, 13, 19 from left to right in 3 cultures. Scale bar: 5 µm). Red asterisks indicate EB3 comets and blue arrow-heads indicate newly formed branches. c. *top*: Schematic and EM images of axon cross-sections at the corpus callosum. Black dots are microtubules. *bottom*: Distance of microtubules to the axon membrane (n=219, 184, 208 in WT, ABfs/fs, L1-Y1229H accordingly in 2 mice/genotype, P<0.0001 in both WT vs. ABfs/fs and WT vs. L1-Y1229H; Scale bar =100 nm). d. Schematic of axon branch formation in ABe37fs/fs and L1CAM Y1229H neurons in comparison to WT. Blue=dynamic microtubules, magenta= hyperdynamic microtubules, red=giant ankB, black= L1CAM. (Mean ±s.e.m, a. bar 1-3, b, d: one-way ANOVA followed by Dunnett’s multiple comparisons test; a. bar 4-8 are t-test. * P < 0.05; ** P < 0.01; *** P < 0.001, **** P < 0.0001).

Collateral axon branching requires microtubules to invade and stabilize nascent actin-based filapodia^25,26^, suggesting the possibility that the increased branching in ABe37fs/fs neurons may result from altered microtubule dynamics. Live-imaging of fast-growing microtubule ends using EB3-tandem tomato (tdTM) as a marker showed increased numbers of microtubules entering filopodia emerging from ABe37fs/fs axons (Fig. 3b). These filipodia, if further stabilized, could lead to an established axonal branch^26^. Further characterization of EB3-tdTM dynamics revealed decreased run length (Extended Data Fig. 3c), indicating that microtubules are hyper-dynamic in ABe37fs/fs neurons. To determine if hyper-dynamic microtubules are the cause of increased branching in ABe37fs/fs neurons, we acutely stabilized microtubules with 0.5nM taxol. Strikingly, taxol-treatment resulted in a marked reduction in axon branches (Fig. 3a).

In order to examine the effect of the giant ankB mutation on the subcellular distribution of microtubules *in vivo*, we imaged cross-sections of unmyelinated axons in the corpus callosum from brains of PND28 mice using transmission electron microscopy (Fig. 3c). Control axons exhibited a cortical microtubule population centered around 20-30 nm from the plasma membrane. In contrast, microtubules in ABe37fs/fs axons were arranged in a broader, lower amplitude peak shifted to 40-50 nm from the membrane (Fig. 3c). Notably, there were no changes in the overall numbers of microtubules per axon, indicating that the cortical population was displaced rather than disassembled (Extended Data Fig. 3d). Taken together these results show that mutation of giant ankB causes increased axonal branching due to hyper-dynamic microtubules and loss of the cortical microtubule population (Fig. 3d).

## A L1CAM/giant ankB/cortical microtubule pathway

We next addressed how giant ankB coordinates cortical microtubules by evaluating the hypothesis that giant ankB is recruited to the plasma membrane through binding to L1CAM. Ankyrins bind to L1CAM through their ANK repeats, which are present in both the 220kDa and giant ankB^27^. However, we found using a proximity ligation assay that L1CAM exclusively interacts *in vivo* with giant ankB but not with 220kDa ankB (Fig. 4b). We observed a strong proximity signal along axon tracts of the corpus callosum in WT mice using one antibody specific for the cytoplasmic region of L1CAM and another antibody recognizing both the 220 kDa ankB and giant ankB (Fig. 4a). This proximity signal was lost in ABe37f/f Nestin-Cre+ mice, which lack giant ankB but still express 220 kDa ankB (Fig. 4b). The signal was also lost in L1CAM Y1229H mice, where L1CAM lacks ankyrin-binding activity (Fig. 4b). We next determined if 290 kDa ankB expressed in ABe37fs/fs mice interacts with L1CAM by performing proximity ligation assays using an antibody recognizing the 290 kDa polypeptide (Fig 1). Remarkably, axons from ABe37fs/fs mice exhibited a marked reduction in the proximity ligation signal, even though 290 kDa ankB retains an L1CAM binding-site (Fig 4b bottom panels). Possible explanations for loss of this interaction are that the 290 kDa polypeptide is expressed at reduced levels and that it also exhibits loss of axonal polarization (Extended Data Fig. 4b).

**Fig. 4.**
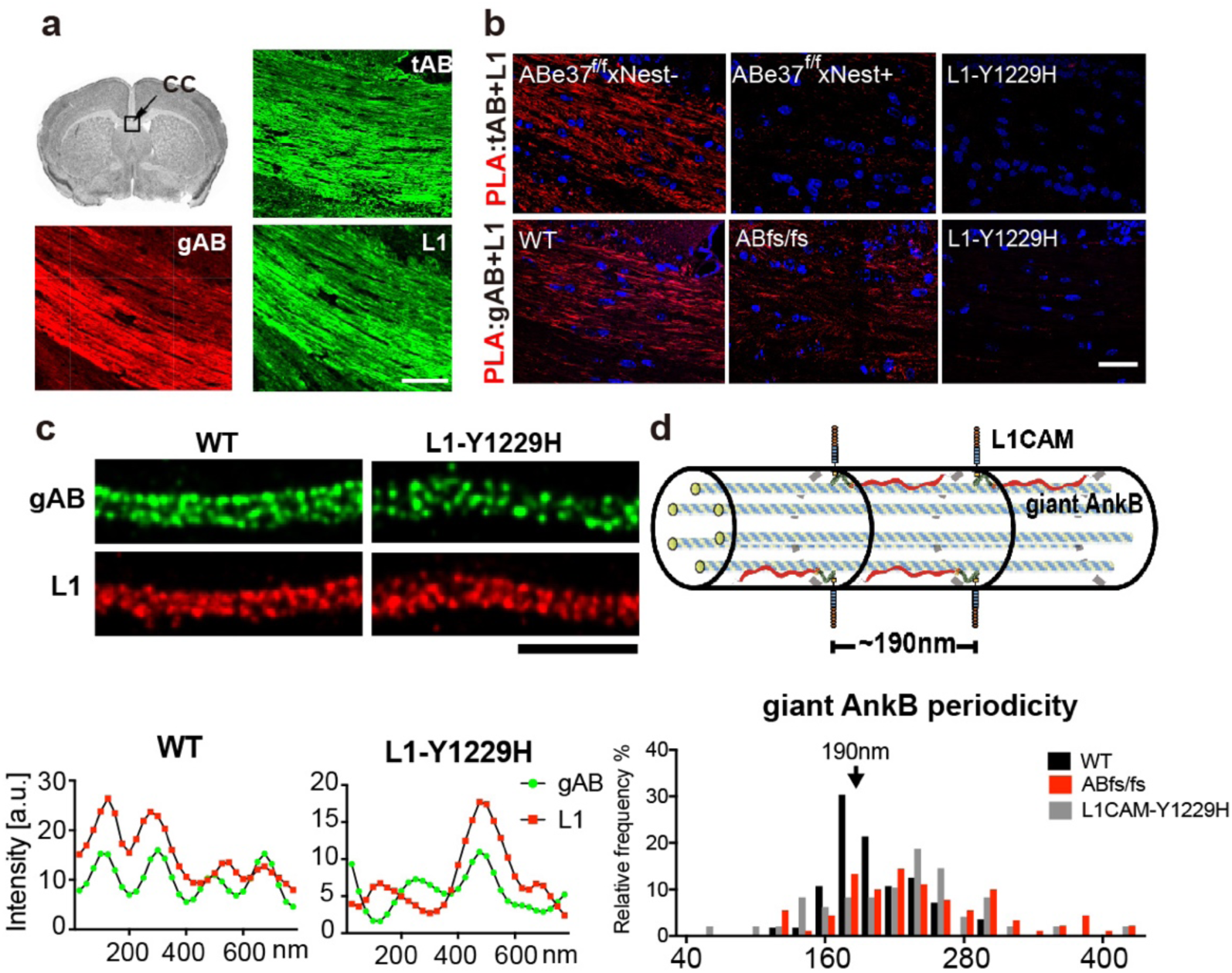
Giant ankB complexes with L1CAM in a periodic pattern in axons. a. Immunostaining for total ankB, giant ankB and L1CAM in corpus callosum of mouse brain coronal sections. b. Proximity signal (red) of the corpus callosal region of WT and indicated mutant mice with Ig against L1CAM and either total ankB (*top*) or giant ankB (*bottom*). Nuclei are in blue (scale bar: 50 µm). c. STED images of giant ankB (red) L1CAM (green) on axons. The staining intensity along the membrane is plotted and the giant ankB periodicity is measured (n= 18, 39, 21 in WT, ABfs/fs, L1CAM-Y1229H. One-way ANOVA followed by Dunnett’s multiple comparisons test; P=0.0113 in WT vs. ABfs/fs, P=0.0003 in WT vs. L1-Y1229H. Scale bar: 2µm). d. Schematic of giant ankB/L1CAM membrane complexes coordinating intracellular microtubules in an axon.

We used STED imaging to resolve the subcellular localization of giant ankB and L1CAM on the plasma membrane of axons in cultured hippocampal neurons (Fig. 4c). Interestingly, axonal giant ankB and L1CAM molecules were regularly spaced at a 190nm periodicity, indicating that they were likely patterned by spectrin-actin rings (Fig. 4c)^28^. The periodicity of giant ankB and L1CAM was lost in both ABe37fs/fs and L1CAM Y1229H cultured neurons, where both proteins exhibited a wider and irregular distribution of spacing (Fig. 4c).

These data indicate that L1CAM and giant ankB are in a complex concentrated in periodic axonal microdomains and suggest that loss of this complex contributes to the axonal phenotypes seen in ABe37fs neurons. We therefore determined the consequences of the impaired L1CAM association with giant ankB on axonal branching and microtubule behavior using neurons derived from L1CAM Y1229H mutant mice. L1CAM Y1229H neurons exhibit increased axon branching (Fig. 3a), increased EB3-tdTM-labeled non-axial microtubules and decreased running length of EB3 (Fig. 3b, Extended Data Fig. 3c). Moreover, axons in brains of L1CAM Y1229H mice exhibited a similar loss of cortical microtubules as observed in ABe37fs/fs mice (Fig. 3c). Together, these results demonstrate that L1CAM is the principal membrane tether for giant ankB, which in turn coordinates cortical microtubules to repress axonal branching.

Interestingly, a similar mechanism utilizing giant ankyrins to control microtubule position in axons has independently evolved in fruit flies^29^. However, giant exons of *Drosophila* Ank2 and vertebrate ankyrins share no sequence homology and are located at different sites within their transcripts^29,30^. The similarities of mechanism thus represent a striking example of convergent evolution.

## Giant ankB mutation indirectly affects excitatory synapses

We next investigated the effects of ABe37fs mutation on synapse number and function at PND28 (corresponding to adolescence in mice) and adulthood (PND 60). The number of presynaptic puncta identified by synaptophysin immunofluorescence labeling was increased in the somatosensory cortices of heterozygous and homozygous ABe37fs mice at PND28 but returned to normal at PND60 (Fig. 5a). We then utilized Golgi-Cox labeling to visualize dendritic spines of pyramidal neurons in the same region and found an increase in the number of mature spines at PND 28 that also returned to normal by PND60 (Fig. 5b). Giant ankB is absent in both the pre- and post-synaptic regions (Extended Data Fig. 5a); thus, the increase in synapses at PND28 is an indirect consequence of the giant ankB mutation, and likely due to increased axon branching.

**Fig. 5.**
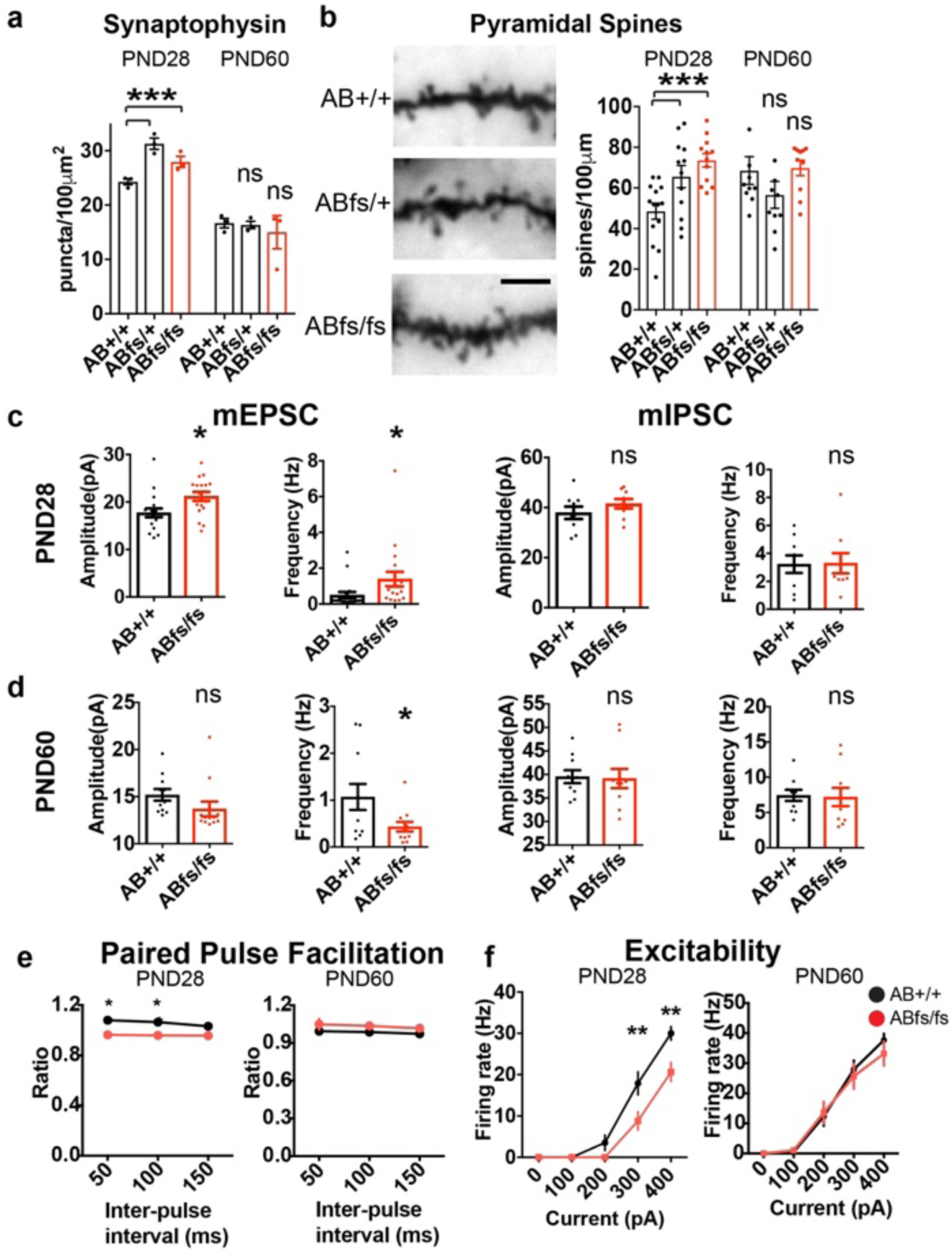
Transient altered synaptic function in PND28 ABe37fs mice. a. Quantification of synaptophysin puncta density in the somatosensory cortex. (N=3 mice/genotype). b. Representative images and quantification of spine densities from apical dendrite branches of pyramidal neurons in the somatosensory cortex. (N=3mice/genotype). c,d. Quantification of mEPSCs and mIPSCs at PND28 and PND60 in acute slices from mice using whole-cell patch clamp recording. Dots represent individual neurons. (N=10mice/genotype). e. Quantification of paired-pulse facilitation generated by an inter-pulse interval (50 to 150ms) in PND28 and PND60 mice (N=8mice/genotype/age). f. Quantification of action potentials generated by direct intracellular current injections in current-clamp recordings in PND28 and PND60 mice (N=8mice/genotype). (Mean ±s.e.m; a,b *** P<0.001, one-way ANOVA, Dunnett’s multiple comparisons test; c,d * P<0.05, t-test; e,f repeated measures two-way ANOVA, Bonferroni posthoc test).

We next evaluated the function of synapses in the ABe37fs mice at PND28 and PND60 using electrophysiology. Whole cell patch clamp recordings from the somatosensory region of acute brain slices at PND28 revealed an increase in frequency and amplitude of mEPSCs in ABe37fs/fs neurons, with no change in mIPSCs (Fig. 5c). These results demonstrate that loss of giant ankB selectively alters the number of excitatory synapses. Due to the increase in mEPSCs, we predicted compensatory synaptic scaling would occur, since mice did not exhibit obvious seizure activity^31^. Indeed, ABe37fs/fs synapses have altered pre-and post-synaptic properties consistent with indirect homeostatic compensation for the increased mEPSCs. ABe37fs/fs neurons showed a decreased paired-pulse ratio, and thus less facilitation of a second EPSC, indicating presynaptic homeostasis (Fig.5e). In addition, we found a significant reduction in neuronal excitability with no change in the resting membrane potential at PND28, as determined by measuring action potentials at fixed levels of current injection (Fig. 5f) (Extended Data Fig. 5b). The reduced excitability is likely a postsynaptic mechanism of homeostasis that compensates at least partially for increased excitatory input.

The changes in synaptic physiology at PND28 were transient, and by PND60 mEPSC amplitude, excitability, and the paired-pulse ratio were all indistinguishable from controls while mEPSC frequency was decreased (Fig. 5a-b, d-f). The transient increase in glutamatergic transmission during juvenile development could facilitate the maturation of ectopic circuitry^32–34^ initially formed by exuberant axon branching. The increased mEPSCs that we observe in PND28 but not PND 60 mice may underlie childhood epileptic seizures observed in some but not all of humans bearing *ANK2* mutation (Case Reports; extended table 1). The restoration of normal synaptic physiological properties at PND60 may result from synaptic pruning and will be important to study in more detail.

## Giant ankB mutant mice exhibit penetrant innate social deficits

We next addressed effects of Abe37fs mutation on behavior. We screened homozygous ABe37fs/fs mice with behavioral tasks evaluating motor, anxiety-like, sensorimotor, cognitive, and social responses. These mutants displayed normal motor abilities on the accelerating rotorod assay, gait, and grip-strength assays (Extended Data Fig. 6a, Extended Data Table 2), with slightly decreased activity in the open field (Extended Data Fig. 6a). No exceptional stereotyped behaviors were noted in the water-spray test or in a marble burying assay (Extended Data Fig. 6b). Abe37fs/fs mice also did not exhibit anxiety-like behaviors, as assessed in the light-dark emergence test and in analysis of time in the center zone of the open field (Extended Data Fig. 6b). Prepulse inhibition was increased (Extended Data Fig. 6c), suggesting that pre-attentive functions may be enhanced in the ABe37fs/fs mice.

Examination of cognitive functions revealed normal short- and long-term novel object recognition memory, and acquisition performance in the Morris water maze and water T-maze (Extended Data Fig. 6d). Surprisingly, when executive function was evaluated during the reversal phase of the water T-maze test^35^, ABe37fs/fs mice demonstrated enhanced performance, as evidenced by shorter latencies to swim to the new platform and by making fewer errors than the controls (Fig. 6c).

**Fig. 6.**
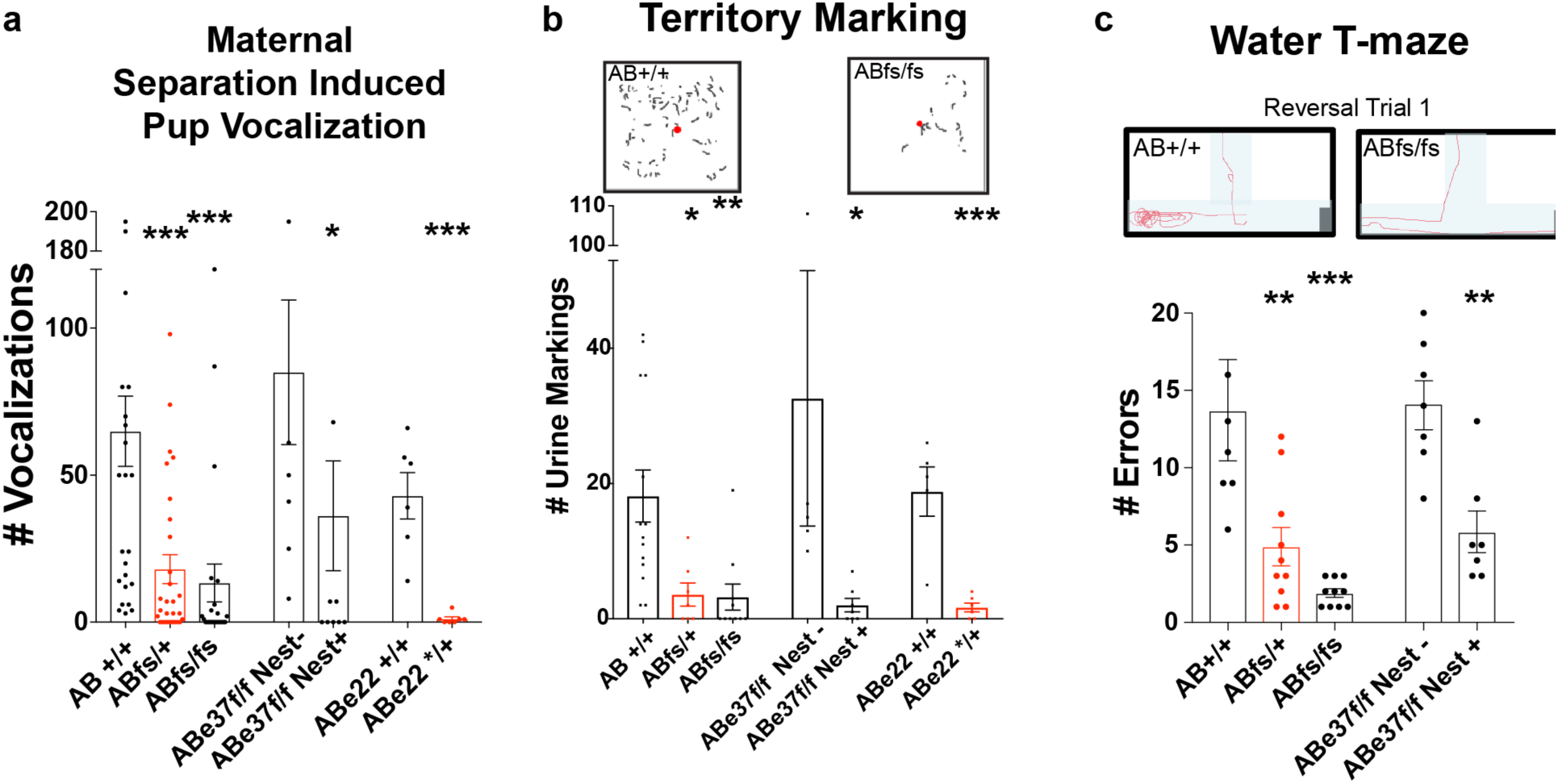
Giant ankB mutant mice exhibit abnormal social responses and increased executive function. a. Quantification of ultrasonic vocalizations from PND7 mice, induced by maternal separation. n=27,29,23,8,13,6,7 pups (from left to right). b. Territory marking by adult males in response to estrous female urine. Representative traces from ninhydrin-exposed urine marks from ABe37+/+ and ABe37fs/fs mice with quantification of marks. n=14,7,10,5,7,6,5 mice (from left to right). c. Reversal performance in a water T-maze: representative traces of the location of the arena in light blue with the hidden platform on the right. Red line shows the trajectory of a mouse starting from the top center with quantification of errors on reversal trial day 1. n=7,10,10,7,7 mice (from left to right). Dots represent individual mice. (Mean ±s.e.m. *P<0.05, **P<0.01, ***P<0.001, two-way ANOVA, Dunnett’s multiple comparisons test for fs, t-test for others).

Despite this superior performance in executive function, innate social behaviors in ABe37fs/fs mice were significantly impaired in both adult males and young pups. In a pheromone-induced urine territory-marking assay, where a sexually-primed male marks his territory by tail-marking urine around an estrous female urine spot, ABe37fs/fs males made small or no territory markings in response to the female urine (Fig. 6b) with no obvious deficits in olfaction (Extended Data Fig. 6c). In a maternal deprivation-induced ultrasonic vocalization (USV) assay, ABe37fs/fs PND7 pups emitted fewer USVs compared to control pups (Fig. 6a). Similar phenotypes of altered pup USVs are reported in D2 dopamine receptor null mice^36^, tryptophan hydroxylase 2 null mice^37^, and *shank3* null mice^38^. Both decreased pup USVs and territory marking are noted in *shank1* null mice^39^. It should be emphasized that heterozygous ABe37fs/+ mice exhibited the same behavioral differences as homozygous mutants (Fig. 6a-b). Hence, these deficiencies are penetrant in mice bearing mutation in one copy of the *ANK2* gene.

To determine if the social deficits and superior executive function were due to loss of giant ankB function, we examined two other mouse strains with giant ankB mutations: ABe37f/f Nestin-Cre+ (ABe37floxed) mice where approximately 90 percent of giant ankB is lost, and heterozygous ABe22*/+ mice [models for human R895* (c.2683C>T), R990* (c.2968C>T) *de novo* nonsense mutations associated with ASD] which have a stop codon inserted into exon 22 producing nonsense-mediated decay and a 50% reduction in both the 220kDa and giant ankB polypeptides. All 3 strains of mice, including ABe37fs/+ and ABe22*/+ animals, had equivalent deficits in both USV and territory marking tests (Fig. 6a-b). Of note, the enhanced cognitive flexibility in the reversal phase of the water T-maze was only reproduced in the 2 strains with targeted deficiency/truncation of giant ankB (i.e., ABe37fs and ABe37floxed) (Fig. 6c). This superior response was not observed in ABe22*/+ mice that have reductions both giant ankB and 220kDa ankB (Fig. 6c). Collectively, these results indicate that giant ankB-deficiency results in dominantly inherited impairment in social and communication behaviors combined with superior executive functions.

## DISCUSSION

We identify giant ankB as a functional neurospecific target for neurodevelopmental mutations in *ANK2* and reproduce penetrant behavioral differences in three mouse models with mutated/deficient giant ankB. We find stochastic as well as regional stereotypical increases in structural connectivity in the cerebral cortices of giant ankB mutant mice using DTI. We deduce a new cellular mechanism dependent on giant ankB that limits ectopic axon branching and is a candidate to explain the aberrant structural connectivity. We find that giant ankB is recruited to periodic plasma membrane domains by L1CAM, where it coordinates cortical microtubules and prevents microtubule entry into and subsequent stabilization of nascent axonal filopodia. This pathway is impaired in mice with mutated giant ankB as well as in L1CAM Y1229H mutant mice where L1CAM lacks ankyrin-binding activity. We also find giant ankB mutation results in increased synapse number and excitatory synapse function at PND28 (corresponding to early adolescence in mice) that is lost in adults, and likely is secondary to axon branching. Thus, stochastic increase in structural connectivity is a candidate to produce the heterogeneous clinical phenotypes seen in *ANK2* variant humans (Fig. 1f,g; Extended Data Table 1).

Increased axon branching has been reported in *PTEN* deficient mice (a high-confidence syndromic ASD-susceptibility gene)^40^. However, giant ankB is unique among the previously elucidated monogenic causes of autism because it is expressed only in neurons (Fig. 1e; Extended Data Fig. 1a). Interestingly, other high-confidence ASD-associated genes including *DYRK1A* (a microtubule kinase)^41^, and *KATNA1* (a microtubule-severing protein)^42^, also affect axonal microtubules. Because of the range of cell types in which these genes are expressed, their mutations are likely to result in multisystem phenotypes in addition to neurodevelopmental abnormalities.

Giant ankB mutation likely affects functional connectivity through mechanisms in addition to axon branching. For example, periodic L1CAM domains were recently reported to be aligned between axons^43^. Loss of this level of axon organization may also occur in giant ankB mutants and could further contribute to loss of axon bundling and ectopic axon tracts. Giant ankB-dependent organization of L1CAM could also be involved in L1CAM interaction with neuropilin-1 which is required for sema3a dependent axon guidance^44^. Giant ankB expression is restricted to pre-myelinated and unmyelinated axons and is increased in myelin-deficient Shiverer mice^16^. It is possible that loss of giant ankB could result in earlier/enhanced myelination, which would affect the speed of action potential conduction and establishment of neural circuits.

Although giant ankB is the direct target for mutation in our mouse models, expression of many genes likely are altered in compensatory homeostatic responses. For example, cultured neurons from WT and ABe37fsfs mice exhibited significant differences in expression of over 400 genes. GO analysis indicates the most significant changes in several synaptic and axon groups in addition to a custom ankyrin-related GO term (Extended Data Table 4, Extended Data Table 5). These alterations in large part may be homeostatic changes allowing the mice to survive loss of a protein conserved throughout vertebrate but may also contribute to abnormal connectivity and behavior. These results illustrate how even a simple monogenic perturbation can have complex consequences and emphasize the challenges in deducing primary molecular mechanisms from gene expression data.

Giant ankB mutant mice exhibit both stochastic and stereotypic alterations in connectivity. The stereotypic connectivity changes are most likely underlie the penetrant phenotypes observed in all three mouse models (Fig. 6). The human correlates for these differences in mouse behavior remain to be determined. Conversely, the stochastic alterations in connectivity observed in mice are a likely candidate to explain the heterogeneous clinical phenotypes seen in *ANK2* variant humans (Fig. 1f,g; Extended Data Table 1). Direct demonstration of divergent behavior between individual mice with the same genotype will require sequential determination of individual structural connectivity and performance for a large number of animals.

The heterogeneous neurodevelopmental phenotypes associated with *ANK2* mutation in humans provides major challenges for traditional approaches to clinical categorization. However, the increasing application whole exome sequencing promises to open a new era of personalized molecular diagnosis. We predict that two distinct classifications within *ANK2* patient variants will resolve: one where the mutation only affects exon 37 which would produce a neurological phenotype, and second where the mutation affects both the 220kDa ankyrin as well as the giant ankB which would have in addition long QT cardiac abnormalities and metabolic syndrome^9–12^. *ANK2* variants found by exome sequencing can be functionally tested in our molecular replacement assay which evaluates axon branching (Fig. 3a). Based on the current exome variant server, a database of over 6000 individuals of both African American and European American ethnicity, there is a single frameshift mutation in *ANK2* exon 37 (Q2018fs*7) carried by 17 individuals^45^. Moreover, 11.7% of the population harbor a possibly to probably damaging polyphen scored missense mutation in exon 37 of *ANK2* (Extended Data Fig. 7, Extended Data Table 6). In particular, 2 variants are present at greater than 3% of the population indicating that they could be under positive selection. Indeed, ASD has been associated with positive selection due to the involvement of neurogenesis and cognitive ability genes^46^. Individuals heterozygous for *ANK2* mutation may experience benefits as well as costs, while much rarer compound heterozygous individuals could suffer a more severe phenotype. Thus, genetic classification is necessary to properly identify neurodevelopmental and other conditions caused by *ANK2* mutations so that treatments, when developed, will appropriately target the cause of the phenotype.

In summary, giant ankB mutant mice provide a neurospecific model for increased ectopic connectivity which we propose is a shared pathophysiology of axon-expressed genes in neurodiverse/divergent individuals and may have diverse consequences in clinical presentation due to the stochastic alterations to connectivity. Indeed, other neurodiversity related conditions, synesthesia and conduct disorder, are associated with increased structural connectivity^47,48^. More generally, stochastic increases in connectivity due to branching of axons may provide novel substrates for neurodiversity with benefits in addition to limitations.

## Case reports

### Patient 1

Patient 1 is a 16-year old Caucasian male with a complex medical history of intellectual disability, epilepsy, and action tremor. He is the product of an uncomplicated pregnancy. At delivery, he is reported to have had bilateral hip and knee contractures and was very tremulous. At 6 weeks of age, he was reported to have his first seizure. Parents report he had 15 seizures that day. He was placed on phenobarbital for seizure management. He continued with this until 3 years of age when he switched to valproic acid. During this interim, he had global developmental delays. He rolled over at 6-9 months, crawled at 12 months, walked at 18 months, and did not talk until 3 years after being taken off phenobarbital. He underwent a left orbitofrontal topectomy at 7 years of age and started on Trileptal. Seizures have been well controlled since surgery on Trileptal. He was attempted to be weaned off seizure medications a few years ago but was unsuccessful. He also has an essential action tremor. He is reported to constantly have tremors; parents have noticed them in his cheeks, eyes, jaw, and tongue. They significantly impair his fine motor skills. He is currently on Inderal after previously being weaned off oxcarbazepine. He is currently a junior in special education high school classes. He has always been in special education classes and will transition to a life skills school after high school graduation. Parents report most recent IQ testing indicates he functions at an 8 to 11-year-old level. Parents deny any behavior concerns. He does have a history of adverse reactions to auditory stimuli and he typically must wear headphones or ear plugs in school and new environments. He is reported to be social and interactive at home. He responds to questions and engages in conversation when prompted. He is also reported to have sleep disturbance. Most recent sleep study indicates he never reaches rapid eye movements. He sleeps for 1-3 hours at a time and has daytime sleepiness; he sleeps for a few hours at school each day. He is non-dysmorphic. Brain MRI in 2009 and 2016 were both non-revealing. Chromosome microarray and Fragile X testing from Baylor Genetics were normal. Clinical trio exome sequencing was subsequently sent and a *de novo* c.4341delC (p.L1448*) variant in the *ANK2* gene was detected.

### Patient 2

Patient 2 is a 7-month-old female who presented with seizure and hypomagnesemia on day of life 8. She is the product of a full term, uncomplicated pregnancy with a routine neonatal course. On day of life 8, she was noted to have left upper and lower extremity twitching intermittently, as well as neck and right shoulder twitching. There was no cyanosis or eye deviation. CBC, chemistry and liver panel were unremarkable except for hypomagnesemia 1.1 (normal range 1.6-2.4 mg/dl). Acylcarnitine analysis, plasma amino acids, cerebrospinal fluid amino acid, urine organic acids and urine purine panel were all normal. Plasma pipecolic acid was elevated (3.4 uM/L, normal <3.0) while cerebrospinal fluid pipecolic acid was normal. Serum calcium and urine Ca/Cr ratio were normal. Head CT, brain MRI and MRI spectroscopy, and renal ultrasound were normal. She took Keppra, phenobarbital, trileptal, and pyridoxine for seizure management. She started weaning off seizure medications with good success at 6 months of age. Her hypomagnesemia resolved around 2 months of age. Chromosome microarray and trio exome sequencing was completed, and a c.8447delG (p.G2816Afs*46) paternally inherited variant in the *ANK2* gene was detected.

Patient 2’s father is 32-year-old male. At 17 days of life, he had seizure-like episodes and presented to a hospital in Turkey. He was provided medication for seizure management. Seizure-like episodes continued for 2-6 months and spontaneously resolved. Episodes have not recurred. He is reported to have developmental delay, specifically he did not walk until 3 years and was reported to be noticeably small in early childhood. He did well in school and has a bachelor’s degree in chemistry and master’s degree in education. He currently works as a testing coordinator at a school. He suffers from chronic pain in his feet but has not had recurrence of seizure episodes.

### Methods

All animal experiments were conducted in accordance with the Duke University Institutional Animal Care and Use Committee guidelines (IACUC Protocol Numbers A167-12-06, A149-15-05, A083-18-04, A153-17-06). Protocol for the mouse behavior core under which all mouse behavioral studies were conducted was A153-17-06.

Written informed consent from patients were obtained in accordance with protocol approved by the Institutional Review Board for Baylor College of Medicine.

### Mice

Abe22* mice were previously generated and are backcrossed to C57BL/6J for >20 generations as previously reported by our lab^19^.

L1CAM Y1229H mice were previously reported^21^. A new qPCR based genotyping strategy was used. A 121-bp amplicon was generated using Forward: ACGTCCAGTTCAATGAGGAT; and Reverse: GGGCTACTGCAGGATTGATAG primers. Alleles were identified by probes (WT: FAM/CA+G+T+ACA+G+TGGC/Iowa Black FQ; Mutant: HEX/CA+G+CA+C+T+CGG/Iowa Black FQ) (Integrated DNA Tech, Coralville, IA).

C57BL6/J mice to backcross strains were obtained from Jackson Laboratories (JAX 000664; Bar Harbor, ME).

### Antibodies for immunohistochemistry

The rabbit anti-total ankyrin-B was previously generated in our lab (1:1000)^49^. Rabbit anti-giant ankB was generated as described below (1 microgram/ml). Mouse anti-L1CAM antibody (1:100 for cultured cells, 1:500 for paraffin sections) antibody was from Abcam (AB24345; Cambridge, MA). Chicken anti-MAP2 (1:1000) was from Abcam (AB5392). Mouse anti-synaptophysin (1:1000) was from Sigma (S5768; St. Louis, MO). Mouse anti-Homer (1:500) was from Synaptic Systems, (160 011; Goettingen, Germany).

### DNA constructs

Full length 440kDa Ankyrin-B cDNA

440kDa Ankyrin-B cDNA was based on a human annotated giant ankyrin-B (ENST00000264366.10). Human Exon37 plus flanking sequence of the cDNA from BstZ17I to BstB1 was synthesized (Genewiz; South Plainfield, NJ) and inserted into the existing human 220kDa ankyrin-B cDNA vector using restriction enzyme digestion. The cDNA of giant AnkB was then cloned into the multiple cloning site of a pre-constructed vector under the ß-actin promotor and C-terminal Halo-tag sequence (gift from Dr. Gary Banker).

Mutations were generated using the Quik Change Mutagenesis XL Kit (Agilent, Santa Clara, CA) for point mutations and the InFusion Cloning Kit (Takara Bio, Mountain View, CA) for frameshifts. After the mutation was confirmed by sequencing, a region flanking the mutation was cut by unique restriction enzymes and ligated into the pAB His 440AnkB-Halo vector.

EB3-tdTomato was a gift from Dr. Erik Dent (Plasmid #50708; Addgene, Cambridge, MA).

pCAG-GFP was a gift from Dr. Gary Banker.

### Sequence comparisons

G1KL59 (Anole), Q8C8R3 (Mouse), Q01484 (Human) protein sequences were aligned by ClustalW2.

### Human clinical exome sequencing and analysis

DNA extracted from peripheral blood was analyzed by clinical exome sequencing at Baylor Genetics Laboratories as previously described ^50,51^. Briefly, genomic DNA samples were fragmented, ligated to Illumina multiplexing paired-end adaptors, amplified with indexes added, and hybridized to exome capture reagent (Roche NimbleGen; Madison, WI). Paired-end sequencing (2 × 100 bp) was performed on the Illumina HiSeq 2500 platform with a mean sequence coverage of ~ 120×, with ~ 97% of the target bases having at least 20× coverage. All samples were concurrently analyzed by Illumina HumanOmni1-Quad or HumanExome-12 v1 array for quality-control and detection of large copy-number variants (CNVs), absence of heterozygosity (AOH), and uniparental disomy (UPD). Exome data were interpreted according to ACMG guidelines and variant interpretation guidelines of Baylor Genetics as previously described ^52,53^.

### Generation of Abe37fs mice

We generated a knock-in mouse to mimic the *de novo* human variant at amino acid 2608 of human *ANK2*. A guide RNA (ATAGTCAGCATCGGGGTCCG AGG) was designed to create a CRISPR-induced indel in a region close to human 2608. The gRNA along with cas9 were microinjected into C57BL6/J embryos. Chimera mice were screened for mutation at the guide sequence. Germline mice were generated by breeding chimeras to C57BL6/J mice. After germline transmission, mice were genotyped by qPCR. A 107bp amplicon was generated (Forward: GAGCAAGAAGCAAAGCAGAAA; Reverse: TCGTCATTCACTTCTGCTGAATA). LNA probes were used to distinguish between WT (FAM/AG+C+C+TCG+GAC/Iowa Black FQ) and fs alleles (HEX/AG+C+G+TC+G+ACC/Iowa Black FQ) (Integrated DNA Technologies).

### Generation of Abe37floxed mice

We generated a conditional knock-out mouse where exon 37 of the *Ank2* gene was deleted under the control of Cre-recombinase. Using BAC recombineering, exon 37 was flanked by *LoxP* sites and a neomycin-resistance cassette that was flanked by *FRT* sites was inserted between the end of exon 37 and the downstream *LoxP* site. The construct was linearized and electroporated into 129S6/SvEvTac-derived TL1 embryonic stem cells by electroporation. The cells were placed under antibiotic selection, and surviving clones were screened to confirm both 5’ and 3’ modifications. Cells were injected into C57BL/6NHsd blastocysts and chimeric animals were obtained and bred to C57BL/6J mice to produce heterozygous animals through germline transmission. Animals were crossed to Flp-recombinase expressing mice to remove the Neo cassette. Animals were then backcrossed to C57BL/6J mice for >8 generations before behavioral testing. Exon 37 was removed from neuronal and glial precursors by crossing Abe37floxed mice with Nestin-Cre recombinase mice (B6.Cg-Tg(Nes-cre)1Kln/J; JAX 003771). Mice were genotyped using the forward primer: CTGTAATATCTGGACAATCTTGAG and reverse primer: AAACAACACAGCGTCTTTTTCCTG. WT genotyping results in a 293bp product and floxed gave a 402bp product.

### Generation of giant ankyrin-B specific antibody

The rabbit anti–giant ankyrin B antibody was generated in our lab. Five peptides (10mg each) with addition of N-terminal lysine residues to facilitate Schiff base chemistry from ankB residues 1443-1620 were synthesized (Thermo Fisher Scientific, Waltham, MA). The sequences for these polypeptides were: KSHLVNEVPVLASPDLLSE, KAAEEEPGEPFEIVERV, KVNEILRSGTCTRDESS, EEEWVIVSDEEIEEARQK, GLVNYLTDDLNTCVPLPK. The peptides were dissolved in 50mM sodium phosphate buffer (pH7.4) to 10mg/ml, mixed, and coupled overnight with glutaraldehyde-activated rabbit serum albumin (RSA, Sigma Aldrich), (5mg/ml; reacted 60 min/24 degrees with glutaraldehyde (EM grade, 1% final, Sigma Aldrich) followed by overnight dialysis to remove free glutaraldehyde). Peptide-RSA conjugates were mixed 1:1 with Freund’s complete (first immunization) or Freund’s incomplete (subsequent immunizations, Sigma Aldrich) adjuvant and 0.5 ml was injected subcutaneously at multiple sites in 4 New Zealand white rabbits (3-4 months). All rabbit procedures were performed by Duke Laboratory Animal Resources. The final pooled sera after 5 injections was from 3 rabbits that reacted specifically with a 440 kDa polypeptide in WT but not ABe37f/f Nestin-Cre + brain lysates. Sera were diluted 1:1 with 150 mM NaCl, 10 mM NaPO_4_, 1 mM EDTA, 1 mM NaN_3_, 0.2% TritonX-100, and heat-inactivated at 56 degrees for 15 minutes. 70 ml of pooled sera were pre-adsorbed with RSA-sepharose and then adsorbed to the mixed peptides coupled to sepharose (5 ml of sepharose). The column was then washed with 500mM NaCl, 10 mM sodium phosphate buffer, 0.1% TritonX-100 (10X column volume), followed by 2M urea, 0.1 M glycine, 0.1% TritionX-100 (2 column volumes), and finally 150mM NaCl, 10mM sodium phosphate, 1 mM EDTA, 1mM NaN_3_ until A_280_ was below 0.01. Finally, antibodies were eluted with 4 M MgCl_2_ and dialyzed into antibody storage buffer [150 mM NaCl, 10 mM sodium phosphate buffer, 1 mM EDTA, 1 mM NaN_3_, and 50% (vol/vol) glycerol]. The specificity of purified final antibody was tested in immunoblots, cell staining and brain paraffin section staining of brain tissue from wild type mice with ABe37f/f Nestin-Cre + mice as a negative control. The Ig concentration for paraffin brain sections and cultured hippocampal neurons was 1µg/ml.

### Quantitative immunoblot and analysis

Mouse brains were dissected, immediately flash frozen in liquid nitrogen and stored at −80°C until ready to use. Brains were homogenized for 30 seconds in 9 volume/weight 65ºC pre-warmed homogenization buffer [8M urea, 5% SDS (wt/vol), 50mM Tris pH 7.4, 5mM EDTA, 5mM N-ethylmelanamide, protease and phosphatase inhibitors] and heated at 65ºC for 15min until the homogenate was cleared. The homogenate was mixed with 5x PAGE buffer [5% SDS (wt/vol), 25% sucrose (wt/vol), 50mM Tris pH 8, 5mM EDTA, bromophenol blue] and heated for 15min at 65℃. Samples were run on a 3.5-17.5% 0.75mm gradient gel in Fairbanks Running Buffer [40mM Tris pH 7.4, 20mM NaAc, 2mM EDTA, 0.2%SDS (wt/vol)]. Gels for in-gel westerns were immediately fixed in 50% isopropanol plus 7% glacial acetic acid for 15min and washed 2x for 10min with dI water. Gels were incubated with primary antibodies (rabbit anti giant ankyrin-B, 1:2000) in 5% BSA (wt/vol) in TBS overnight at 4°C. After washing, gels were incubated with secondary antibodies (goat anti-rabbit 800CW, 1:10,000; Licor 926-32211; Lincoln, NE) for 2 hours. Gels were extensively washed with TBST and the final 2 washes were performed in TBS. Gels were imaged on the Licor Odyssey at 0.37mm custom offset.

Quantification for lower molecular weight proteins (not giant ankyrins) was done with traditional immunoblot where gels were transferred onto nitrocellulose membranes (Biorad 1620115; Hercules, CA) overnight at 0.3A at 4°C. After transfer, efficiency was determined using Ponceau-S stain. Membranes were blocked using 3% non-fat milk (wt/vol) in TBS for 1 h and then incubated with primary antibodies overnight in blocking buffer at 4°C (rabbit anti-total ankyrin-b 1:2000; mouse anti-GAPDH (ThermoFisher, MA5-15738). Membranes were washed with 3x for 10min and incubated with secondary antibodies (goat anti-rabbit 800CW, goat anti-mouse 680RD, Licor 925-68070) for 2 hours. Membranes were washed 3x for 10min with TBST and 2x for 5min in TBS. The membranes were then imaged on the Licor Odyssey at 0.00mm offset. Blots were quantified using the Image Studio Odyssey software to determine the signal of the band. Background was normalized using a similar % acrylamide region of the image. Values were averaged from 3 technical replicates for each biological sample.

### Immunohistochemistry and confocal microscopy

Immunohistochemistry and confocal microscopy were performed as previously described^54^. Mice of specified genotypes were transcardially perfused at PND28 or PND60 with 4% (wt/vol) paraformaldehyde (PFA) dissolved in PBS, 5 mM Na EDTA, 10 mM NaNO_3_ after flushing out blood with PBS, and the brains were immediately dissected out and post-fixed in the same fixation buffer overnight at 4°C. Brains were then processed for paraffin embedding, microtome cutting, deparaffinization, and immunostaining. Images were collected using 40X/NA1.4 oil or 63X/NA1.4 oil objectives on a LSM780 inverted confocal microscope (Zeiss, Oberkochen, Germany). Z-stacks were collected for thick samples and final images were 3D rendered using Fiji^55^.

### Hippocampal neuron culture, transfection and immunostaining

Primary hippocampal neurons were prepared as described previously^56,57^. Briefly, hippocampi were dissected from genotyped postnatal 0-1 day mice, trypsinized, dissociated, and plated onto poly-L-lysine coated 18-mm glass coverslips in pre-conditioned glia feeder dishes (~200,000 neurons/60mm culture dish). Cultures were grown in neurobasal-A medium plus B27 supplement (2%) and GlutaMAX (Thermo Fisher) and maintained at 37°C incubator with a 5% CO_2_. Constructs were transfected into hippocampal neurons after 3 days of culture with Lipofectamine 2000 (Thermo Fisher) and subsequently fixed with 4% wt/vol) PFA with 4% (wt/vol) sucrose in PBS for 15min at 37°C prior to antibody staining. Fixed neurons were washed with PBS, permeabilized with 0.25% TritonX-100 for 5 min, and then blocked with 5% BSA in PBS for 1 hour. The primary antibodies were diluted in blocking buffer and incubated at 4°C overnight. The next day the neurons were washed with PBS, incubated with the appropriate fluorescent secondary antibodies in blocking buffer for 2 h at room temperature. Finally, the neurons were mounted with Prolong Diamond (Thermo Fisher) onto glass slides and allowed to cure for a minimum of 24 h before imaging by confocal microscopy (Zeiss LSM780 inverted confocal; Oberkochen, Germany). Cells expressing constructs with HaloTag were treated with 50 nM Janelia Farm 549 dye^58^ for 10 min and washed for 10 min prior to fixation or imaging.

### Taxol treatment of cultured neurons

Cultured neurons were transfected at DIV3 with pCAG-GFP using Lipofectamine 2000 to visualize cell morphology. After recovering for 12 hours, cells were treated with 0.5nM taxol (T1912; Sigma-Aldrich) or DMSO as a control for 12 hours. Cells were fixed at DIV4 and imaged as described above. The conditions for the dose of taxol and the duration of treatment were first established in wild-type neurons where the final parameters were found not to alter neuronal morphology.

### EB3 comet imaging and quantification

Cultured hippocampal neurons were transfected at 4 days and imaged the next day on the Andor XD Revolution Spinning Disk Confocal microscope (spinning-disk confocal-head: Yokogawa CsuX-1 5000 rpm) equipped with an EMCCD camera. Cells were maintained in live-cell imaging solution (Life Tech). The whole imaging stage and objectives were maintained at 37°C, 5% CO_2_ in an enclosure. A 100x/NA1.4 oil U PlanSApo DIC objective (Olympus; Center Valley, PA) was used to acquire image streams of 40 frames (3s interval). For the EB3 running length and velocity quantification. MetaMorph software (Molecular Devices; San Jose, CA) was used to generate maximum intensity kymographs from movies. A blinded reviewer identified transport events on the kymograph and coordinates were exported to Excel for analysis of EB3 velocity and running length. A minimum of 10 cells from each condition were evaluated, including cells from at least two independent cultures. For the EB3 non-axial stack images, a series of images from 40 time points were built using the Z-projection macro in Fiji. The contrast of the stack images was inverted to best display the trajectory of EB3 comets.

### Proximity ligation assay

Protein proximity (within 40nm) was assessed using the Duolink PLA kit (DUO92101, Sigma-Aldrich). In brain sections, slides were deparaffinized as above and processed as outlined by the commercial manual using rabbit-derived antibodies against all isoforms of ankyrin-B or just for giant ankyrin-B, and with a mouse-derived antibody against L1CAM. In cultured neurons, cells were fixed and permeabilized as described above and then processed as outlined in the commercial manual with the same sets of antibodies. The images were collected using a 63X/NA1.4 oil objective on a LSM780 inverted confocal microscopy (Zeiss).

### Stimulated Emission Depletion super-resolution imaging and quantification

High resolution images were taken by a Leica 3X STED DMi8 motorized inverted microscope with 100X/NA1.4 objectives (Wetzlar, Germany). The secondary antibodies used for STED images were Oregon Green 488 and Alexa Fluor 594 (1:500 dilution) following our standard immunostaining protocol (described in the immunostaining section). The STED images were de-convolved using the default settings of Huygens deconvolution software linked to LAS X for STED. The intensity measurement for antibody staining was performed in Fiji. The giant ankyrin B and L1CAM periodic measurements were quantified by self-written Matlab scripts (available upon request). The measurements were exported to the GraphPad^®^ for the final graphs.

### Golgi-Cox staining

Golgi-Cox staining was performed on brains from PND28 and PND60 mice according to the manufacturer’s suggestions (PK401; FD Neurotech, Columbia, MD). Brains were frozen and stored at −80°C after impregnation with the staining solutions until they were ready for developing and imaging. Brains were sectioned at 100µm on a cryostat (Leica CM3050 S) and sections were placed onto gelatin-coated slides. After development, sections were mounted with Permount (Electron Microscopy Science, Hatfield, PA) and imaged with a 20X/NA0.8 objective on a widefield microscope (Zeiss Axio Imager Z2). Z-stacks were collected in the somatosensory region of the cortex. For synapse counting, a blinded individual counted spines from a secondary branch of an apical dendrite manually in Fiji. The number of spines was divided by the length of the segment counted to calculate the density of spines. For axon branching quantification, Z-stack images was collected by 10X/NA0.45 objective of the same microscope.

### Electron microscopy

ABe37fs/fs, L1CAM Y1229H hemizygous, and control mice were perfused at PND28 with 2% PFA and 2.5% glutaraldehyde in 0.1 M sodium cacodylate buffer (pH 7.4) at room temperature and this was followed by post-fixation in the same buffer at 4°C for up to 10 days before the brains were dissected. Fixed brains were sectioned using a microslicer to generate ~2mm-thick sagittal sections. Matching corpus callosum areas of each tissue were further dissected, fixed overnight, sliced and processed by the conventional method of the Duke Electron Microscope Core. Final images were observed with a transmission electron microscope (JEOL 1230 TEM) at the UNC microscopy services laboratory. Distances between microtubules to the nearest axonal membrane in the unmyelinated axons of each genotype were measured manually in Fiji. For each genotype, at least two biological replicates were examined.

### Behavioral Tests

Four cohorts of mice were tested where both sexes were examined (except in the urine open-field test where only males were examined). Mice were maintained on a 14:10 h light:dark cycle under humidity- and temperature-controlled conditions with *ad libitum* access to water and chow. All testing occurred during the light phase and mice were acclimated to each behavioral room for at least 24 h prior to testing.

### Open-field activity

Animals were placed individually into the open field for 1 h (OmniTech Inc, Columbus, OH). Fusion software (OmniTech Inc) automatically scored all motor activities over the 1 h interval.

### Grip strength

Strength of the front paws and all 4 paws (whole animal) was assessed with a mouse grip-strength meter (San Diego Instruments, San Diego, CA). Mice were allowed to grip the wire grid(s) with the specified paws and were pulled gently until they released the grid(s).

### Gait analyses

Gait analyses were conducted with the TreadScan apparatus (CleverSys, Reston, VA). Stride length and width, foot placement, the coupling ratio of front and rear paws, and time in phases of gait were calculated.

### Rotarod performance

Accelerating rotarod performance (Med-Associates, St. Albans, VT) was assessed from 4 to 40 rpmb, where a trial terminated when the mouse fell from the rod or after 5 min. Performance was evaluated over 4 trails with an inter-trial interval of 30 min.

### Water-spray test

Animals were individually placed into a clean cage and filmed (MediaRecorder2; Noldus Information Technologies, Ashville, NC). Mice were placed into the test-chamber for 10 min and then were lightly misted with water. Mice were observed for an additional 10 min. Grooming behavior was scored with TopScan software (CleverSys) by a blinded observer.

### Marble burying test

Mice were placed into individual cages with 6 cm of bedding and 20 marbles (Floral blue/green glass marbles 5/8in; Darice, Strongsville, OH) arranged in a 4 by 5 array. Testing occurred over 30 min. Videotapes of marble placement were scored as buried, partial buried, moved marbles, and cavities dug with Ethovision (Noldus) by a blinded observer.

### Light-dark emergence test

Mice were placed in the darkened side of a two-chambered apparatus (Med-Associates) and given 5 min to explore the darkened (<1 lux) and lighted (~750 lux) chambers. Infrared diodes tracked the location and activity of the animal.

### Prepulse inhibition (PPI)

Acoustic startle responses were acquired and analyzed as previously described using the SRL-Lab startle response system and software (San Diego Instruments; San Diego, CA)^59^. Mice were habituated in the PPI chambers for 10 min. After this time, mice were randomly exposed to null trials (64 dB background white-noise), pulse-alone trials (a 40msec 120dB white-noise startle stimulus, or prepulse-pulse trial (20msec prepulse stimuli that were 4, 8, or 12dB above the white-noise background followed 100msec later by the 40msec 120dB stimulus). Each mouse was exposed to 42 trials composed of 18 pulse-alone trials, 6 null trials and 6 prepulse-pulse trials at each intensity. The mean startle response (AU) was recorded. PPI was calculated as the ratio of the mean startle response on prepulse-pulse trials to the pulse-only trials subtracted from 1 and multiplied by 100 to be expressed as a percent.

### Novel object recognition test

Novel object recognition memory was conducted to assess short- and long-term memory. Mice were acclimated to the test apparatus for 10 min. Thereafter, they were exposed to 2 identical objects for 10 min. Short-term memory was assessed 30 min later where one of the identical objects was replaced with a novel shape/color object. After 24 h long-term memory was evaluated where one of the previous objects was replaced with a novel object. All objects were 2.5-3cm^2^. Trials were filmed and analyzed using Ethovision XT 9 (Noldus) where object interaction was scored when the nose of the mouse was within 3 cm of the object. Object preference was calculated by taking subtracting the time spent with the familiar object from time spent with the novel object and dividing this number by the total time spent with both objects. A positive score indicates preference for the novel object, a negative score denotes preference for the familiar object, and scores approaching zero signify no preference for either object.

### Morris water maze

Spatial learning and memory were examined in the Morris water maze. Mice were acclimated to water for 4 days prior to testing. One day before testing, mice were placed on a hidden platform in the northeast (NE) quadrant, allowed to freely swim for 20 s, and permitted to return to the platform. In the acquisition phase of the test, mice were administered 4 trials per day (30 s inter-trial interval) for 8 days where they were released at random points in the maze across trials and days. The trial ended after 60 s when the mouse found the platform or was removed from the maze. Probe trials were given on even days where the mouse was allowed to swim for 60 s. Performance on all tests was video-recorded and scored with Ethovision XT 9 (Noldus).

### Water T-maze

Mice were acclimated to swimming over the course of 4 days. On day 1 mice were placed into the maze in 0.1-0.5cm water and on day 2 they were in 1-2cm water. On days 3 and 4 mice were placed on a hidden platform and allowed to swim in 10 cm of water for 15 s and were returned to the platform after this time. After water acclimation, mice were housed in the test room for at least 24 h. After this time, one-half of the mice were trained to swim to a hidden platform in the left arm and the other half to a platform in the right arm of the maze. Mice were given 4 consecutive trials a day with a maximum swim time of 1 min. Incorrect arm choices resulted in the mouse being retained initially in the incorrect arm for 30 s, after which the barrier was removed, and the mouse was allowed to swim to the platform in the correct arm. Animals had to reach a criterion of 3 correct choices over 2 consecutive days to advance to the next phase of testing. Here, the location of the platform was reversed, and no punishments were made for the incorrect arm choice. Reversal trials ended either when the mouse successfully found the platform or after 60 s. All trials were filmed and analyzed by Ethovision XT 9 (Noldus).

### Pup ultrasonic vocalizations

Individual PND7 mice were placed into a cushioned cup in a styrofoam chamber. An externally polarized condenser microphone was placed 15 cm above the cup and pup ultrasonic vocalizations (USVs) were recorded (10-200 kHz) for 90 s (Avisoft Bioacoustics, Glienicke, Germany). Spectrograms were analyzed with automated whistle tracking parameters (Avisoft SASLab Pro) and confirmed by hand-scoring by a blinded observer.

### Male territory-marking test

The urine-marking open field test was run similar to that described^60^. Males were exposed to wild-type estrous female bedding for 3 days prior to testing. The open field was lined with paper and with a small amount of its own home-cage bedding in the corner. The mouse was allowed to habituate to the arena for 10 min. After this time, the mouse was transferred to a clean cage and the arena was cleaned with all bedding removed. A UV light was used to identify any urine marking and all marks were circled in pencil to. Subsequently, 15µl of estrous female urine, pooled from 4 estrous females, was spotted into the center of the paper. The mouse was returned to the arena for 5 min and then removed to its home-cage. The paper was removed, treated with Ninhydrin spray (Sigma), and left to dry overnight. Papers were scanned, and urine markings were counted by a blinded observer.

### Olfaction ability test

Mice were tested for their ability to discriminate an odor. Mice were placed individually in a clean cage without bedding for 10 minutes to acclimate. After acclimation, two cartridges were placed on the far walls of the cage, each containing a cotton swab with either 10ul almond extract or 10ul saline. Mice were recorded for an additional 10 minutes. Preference was calculated based on duration with the almond cartridge minus the saline cartridge divided by the total time spent with either cartridge.

### Magnetic resonance histology and image analysis

Mice were anesthetized and perfused through the left ventricle with 0.9% saline at a rate of 8 ml/min for 5 min. Fixation was performed with a 10% solution of neutral phosphate-buffered formalin containing 10% (50 mM) Gadoteridol (ProHance; Bracco Diagnostics Inc., Monroe Township, NJ) at the same rate for 5 min. The fixed specimens were transferred to a 0.01 M solution of PBS containing 0.5% (2.5 mM) Gadoteridol at 4 °C for 5-7 days to rehydrate the tissue. Extraneous tissue around the cranium was removed prior to imaging, and specimens were placed in MRI-compatible tubes and immersed in perfluoropolyether (Galden Pro, Solvay, NJ) for susceptibility matching.

MRI was performed using a 9.4T, 8.9 cm vertical bore Oxford magnet, with shielded coils, providing gradients up to 2000 mT/m (Resonance Research, Inc., Billerica, MA), controlled by an Agilent Direct Drive Console (Agilent Technologies, Santa Clara, CA), and custom-made solenoid coils. Imaging protocols relied on 3D spin echo diffusion-weighted acquisition, with 31 diffusion sensitization directions, interspersed with 3 non-diffusion weighted scans. To accelerate acquisition, we used compressed sensing with a compression factor of 4^38^. The imaging matrix was 368×184×184 mm; field of view 22×11×11 mm, repetition time TR=90ms; echo time TE=12 ms; diffusion amplitude = 130.67 G/cm, durations were 4ms, separation was 6 ms, b=4000 s/mm^2^; BW=125kHz. Images were reconstructed at 55 µm isotropic resolution. Total imaging scan time per animal was 7 h and 11 min.

Image Analysis was performed using a high performance computing (HPC) pipelines for image reconstruction relying on the Berkeley Advanced Reconstruction Toolbox^61^; and a pipeline for segmentation^62^, regional and voxel-wise statistics using a symmetric brain atlas with 332 regions (https://arxiv.org/pdf/1709.10483) as refined based on definitions previously used^23^.

Diffusion tensor estimation and tractography based connectomes and interhemispheric asymmetry were calculated using DSIStudio (dsi-studio.labsolver.org/)^22^. 200,000 seeds were randomly placed in the mask brain. Fractional anisotropy was thresholded at 0.07 and the angular threshold was 65°. The step size was 0.03 mm and fiber trajectories were smoothed by averaging the propagation direction with 1% of the previous direction. Region specific analysis was conducted with the same parameters but with 20,000 seeds. R (https://www.r-project.org/) and MATLAB (MathWorks, Natick, MA) were used for subsequent analyses. The interhemispheric asymmetry of FA tracts was determined by the absolute difference of paired pixels between the left and right hemisphere normalized by the total track volume (perfect symmetry=0, perfect asymmetry=1). The whole brain analysis is based on tracks generated from seeds placed over all the brain. Cortex analysis is based on tracks seeded in region 1-41. The MATLAB script used to calculate the asymmetry level is available upon request.

### Whole-cell patch clamp recording

For whole-cell patch-clamp recordings, the brain (PND 28, WT: 10, KO: 10; PND60, WT: 10, KO: 10) was removed quickly and placed into an ice-cold solution bubbled with 95% O_2_-5% CO_2_ containing the following (in mM): 194 sucrose, 30 NaCl, 2.5 KCl, 1 MgCl_2_, 26 NaHCO_3_, 1.2 NaH_2_PO_4_, and 10 *D*-glucose. After 5 min, 250 μm coronal slices were cut and placed in a 35.5°C oxygenated artificial cerebrospinal fluid (aCSF) solution containing the following (in mM): 124 NaCl, 2.5 KCl, 2 CaCl_2_, 1 MgCl_2_, 26 NaHCO_3_, 1.2 NaH_2_PO_4_, and 10 *D*-glucose, pH adjusted to 7.4 with HCl and with the osmolality set to ~320 mosM. After 30 min, the slices were left in aCSF at ~22-23°C for at least 30 min before recording.

All recordings were performed with a MultiClamp 700B amplifier (Molecular Devices, San Jose, CA). Signals were filtered at 10 kHz and digitized at 20 kHz with a Digidata 1440A digitizer (Molecular Devices). During the recordings, the slice was maintained under continuous perfusion with aCSF at 28–29 °C with a 2–3 ml/min flow rate. After achieving whole-cell configuration (series resistance < 25 MΩ), we recorded the miniature EPSCs (mEPSCs), miniature IPSCs (mIPSCs), paired pulse ratio, and excitability.

To measure excitability, pipettes (3.5–5 MΩ) contained the following (in mM): 150 K-gluconate, 2 MgCl_2_, 10 HEPES, 1.1 EGTA, 3 Na-ATP, and 0.2 Na-GTP, pH adjusted to 7.2 with KOH with the osmolality set to ~ 315 mosM. Excitability was measured in current clamp mode by injection of current (−400 to 300 pA). Each step was at 100 pA with a duration of 1 s, and the number of spikes for each depolarizing step was counted. To measure miniature excitatory postsynaptic current (mEPSC), pipettes (3.5–5 MΩ) contained the following (in mM): 120 cesium methane sulfonate, 5 NaCl, 10 tetraethylammonium chloride, 10 HEPES, 4 lidocaine N-ethyl bromide, 1.1 EGTA, 4 Mg-ATP, and 0.3 Na-GTP, pH adjusted to 7.2 with CsOH with the osmolality set to ~ 300 mosM. mEPSCs were recorded with 1 µM tetrodotoxin and 50 µM picrotoxin in the bath solution at −70 mV in voltage-clamp mode. Only mEPSCs with amplitudes greater than 10 pA by peak detection software in pCLAMP10 (Molecular Devices) were included in the analysis. Paired pulse ratio was determined by calculating the ratio of the peak amplitude of the second evoked EPSC to that of the first evoked EPSC (50, 100, and 150 ms interstimulus intervals).

To measure miniature inhibitory postsynaptic current (mIPSC), pipettes contained the following (in mM): 110 cesium methane sulfonate, 30 K-gluconate, 0.1 CaCl_2_, 10 HEPES, 1.1 EGTA, 4 Mg-ATP, and 0.3 Mg-GTP, pH adjusted to 7.2 with CsOH with the osmolality set to ~ 300 mosM. mIPSCs were recorded with 1 µM tetrodotoxin and 20 µM APV and 50 µM DNQX in the bath solution at −70 mV. Only mIPSCs with amplitudes greater than 20 pA were included for analysis.

### RNAseq sample preparation and analysis

Cortical neuronal cultures from P0 mice were grown until DIV14. RNA was isolated using RNeasy Mini Kit (Qiagen). RNAseq libraries were prepared using the TruSeq Stranded total RNAseq kit in combination with Ribozero Gold (Illumina; San Diego, CA). Samples were run on Illumina HiSeq4000. Samples had between 49,000,000 and 65,000,000 passed filter clusters with an average quality score of 39.42. RNA-seq data was processed using the TrimGalore toolkit^63^ which employs Cutadapt^64^ to trim low-quality bases and Illumina sequencing adapters from the 3’ end of the reads. Only reads that were 20nt or longer after trimming were kept for further analysis. Reads were mapped to the GRCm38v73 version of the mouse genome and transcriptome^65^ using the STAR RNA-seq alignment tool^66^. Reads were kept for subsequent analysis if they mapped to a single genomic location. Gene counts were compiled using the HTSeq tool^67^. Only genes that had at least 10 reads in any given library were used in subsequent analysis. Normalization and differential expression were carried out using the DESeq2^68^ Bioconductor^69^ package with the R statistical programming environment^70^. The false discovery rate was calculated to control for multiple hypothesis testing. Gene set enrichment analysis^71^ was performed to identify gene ontology terms associated with altered gene expression for each of the comparisons performed.

### General statistical analyses

Statistical analyses and graph generation were performed using Graphpad Prism 7 (La Jolla, CA) unless otherwise noted. Student’s t-test was used for comparisons between two groups and a one-way ANOVA with Tukey’s *post-hoc* or Bonferroni pair-wise comparisons test or Dunnett’s multiple comparison test were used for comparisons of more than two groups. Repeated-measures ANOVA was used for comparisons of the same groups across time. A P<0.05 was considered statistically significant.

## Supporting information

## Supplementary Information

Supplementary Information is linked to the online version of the paper at www.nature.com/nature.

---

**Extended Data Fig. 1a.**
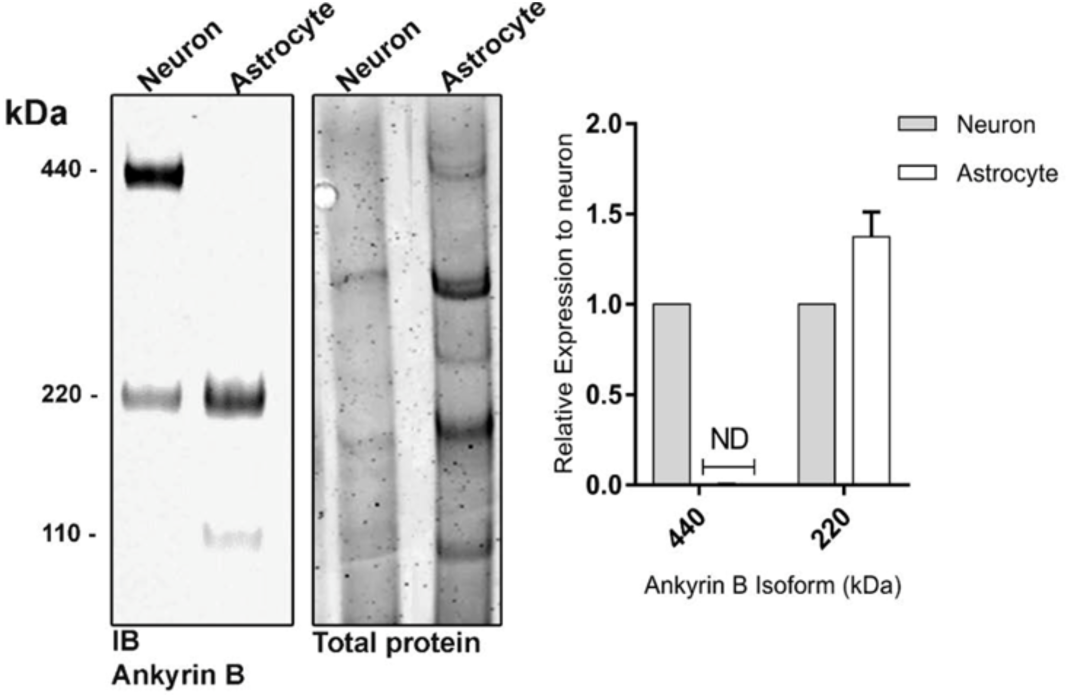
Neuronal specificity of giant ankB. Representative in-gel immunoblot using Ig against the C-terminal region of ankB (recognizes giant ankB, 220kDa and 110kDa ankB) and corresponding colloidal Coomassie blue gel from DIV14 cultured neurons and DIV14 cultured astrocytes with quantification of ankB polypeptide levels. Quantification based on normalization to total protein and neuron level set to 1 AU. n=3 cultures; mean ±s.e.m. t-test.

**Extended Data Fig. 1b.**
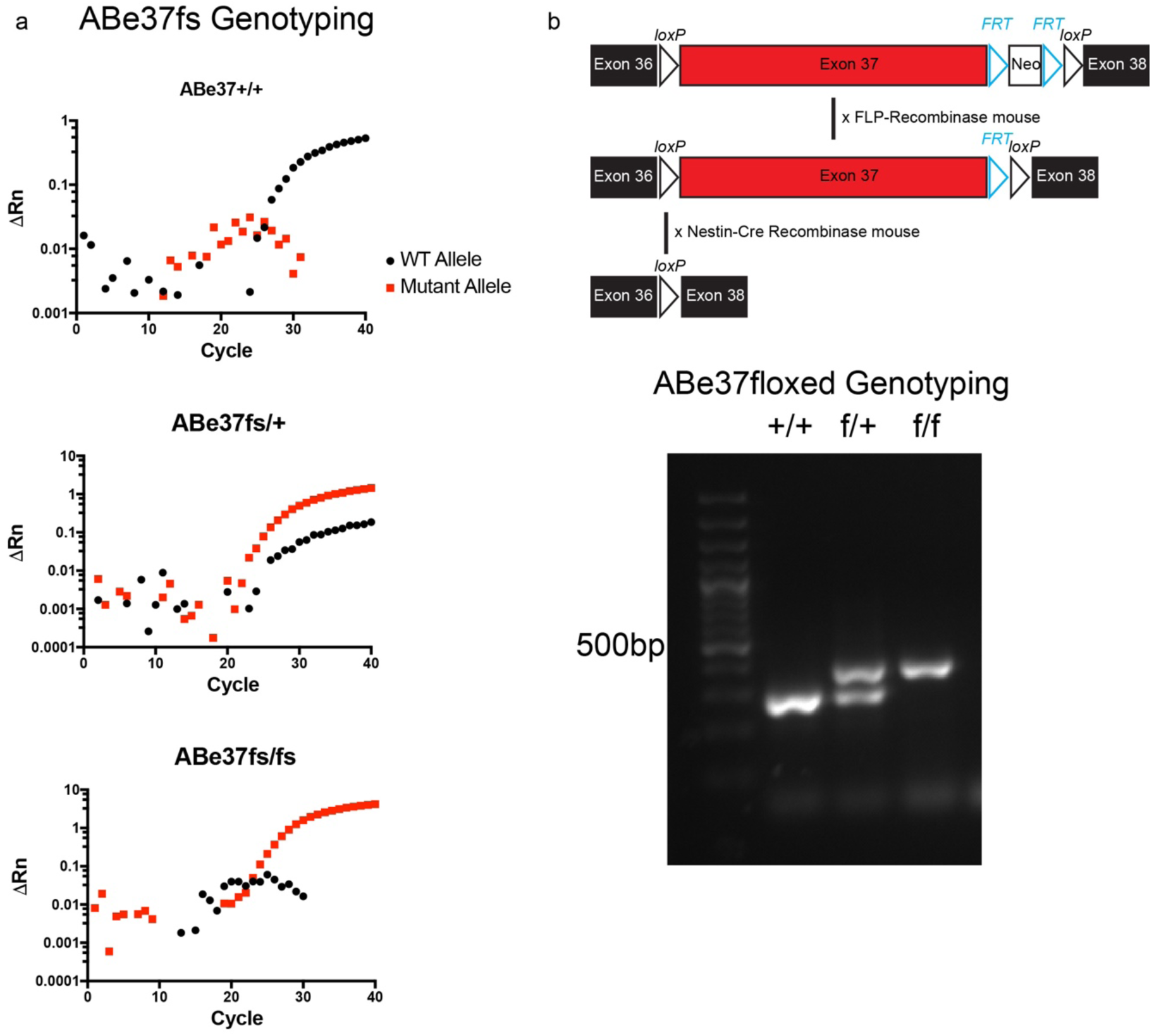
a. qPCR amplification using probes specific for control and frameshift DNA to genotype control, heterozygous, and homozygous ABe37fs mice. b. Scheme of Abe37floxed mouse and DNA gel of PCR amplification for the floxed exon 37 allele in control, heterozygous and homozygous ABe37floxed mice.

**Extended Data Fig. 1c.**
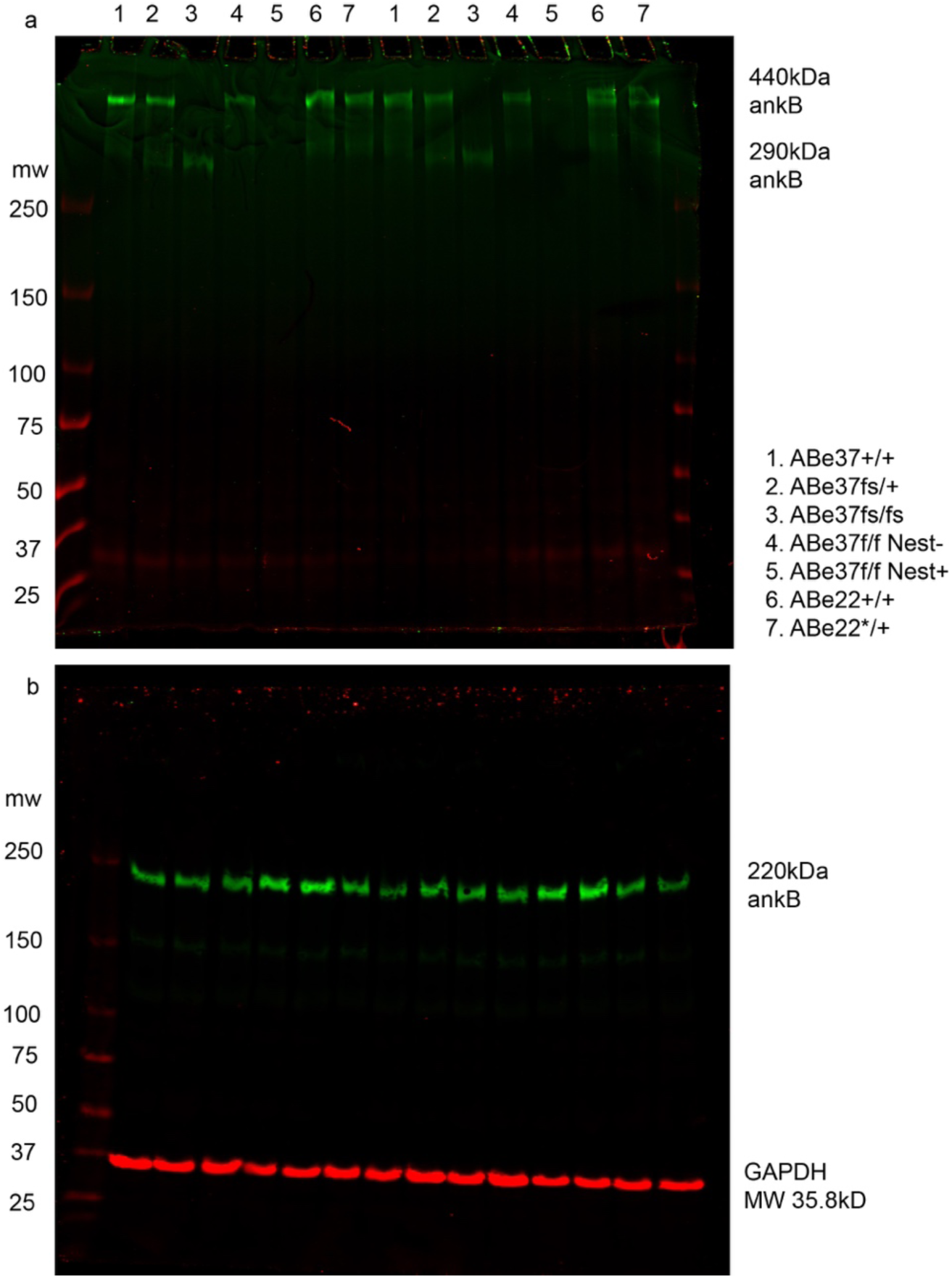
Uncropped immunoblots. a. In-gel Western using rabbit anti-giant ankyrin-B. b. Transferred membrane using rabbit anti-ankyrin-B C-terminal (green) and mouse anti-GAPDH (red). MW based on a protein ladder labeled on the left.

**Extended data Fig. 1d.**
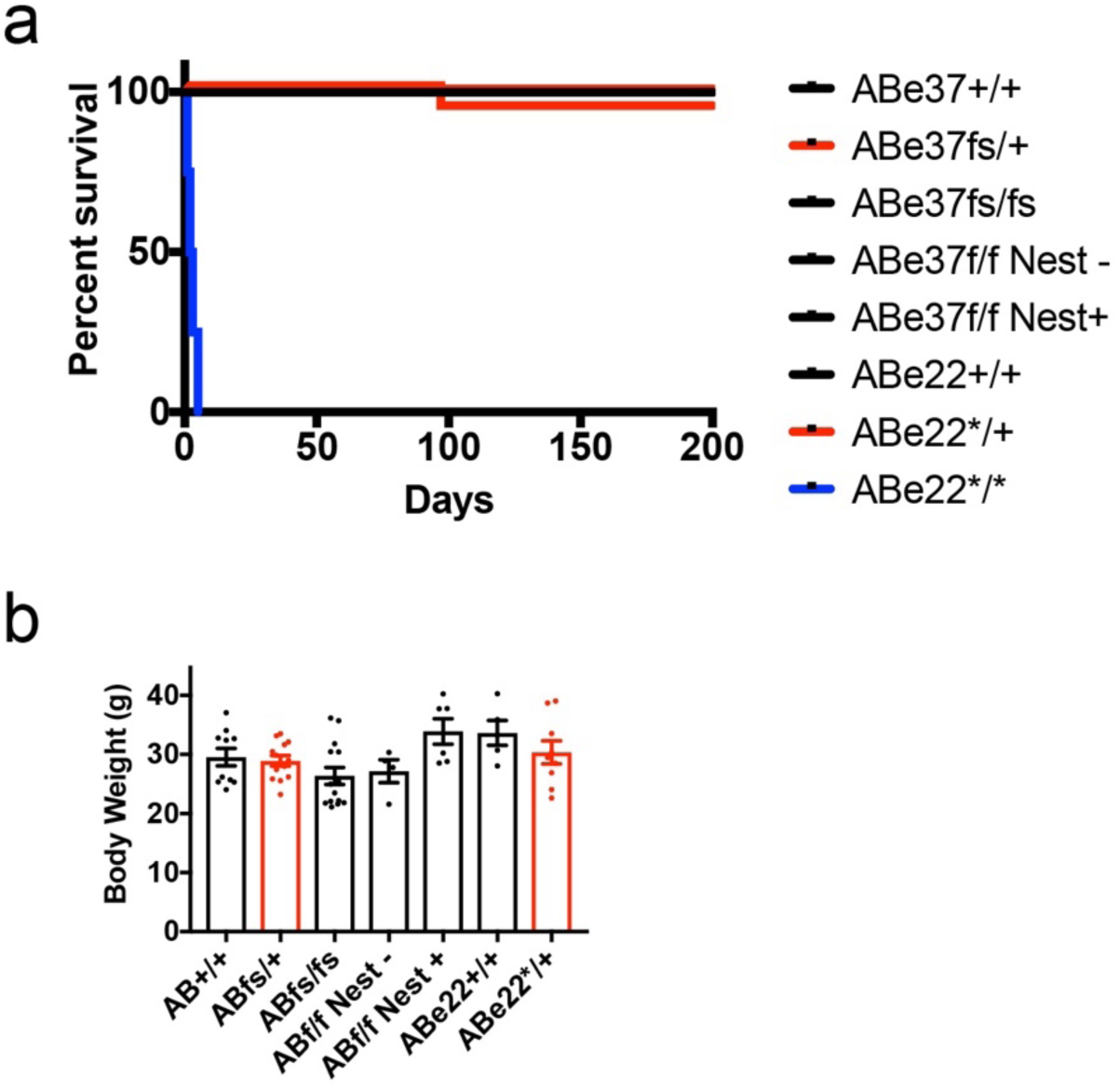
Life span of ANK2 mutant mouse strains and body weight of adult ANK2 mutant mice (a. Log-rank Mantel-Cox test, *P<0.0001 and b. mean ±s.e.m. one-way ANOVA).

**Extended Data Fig 2.**
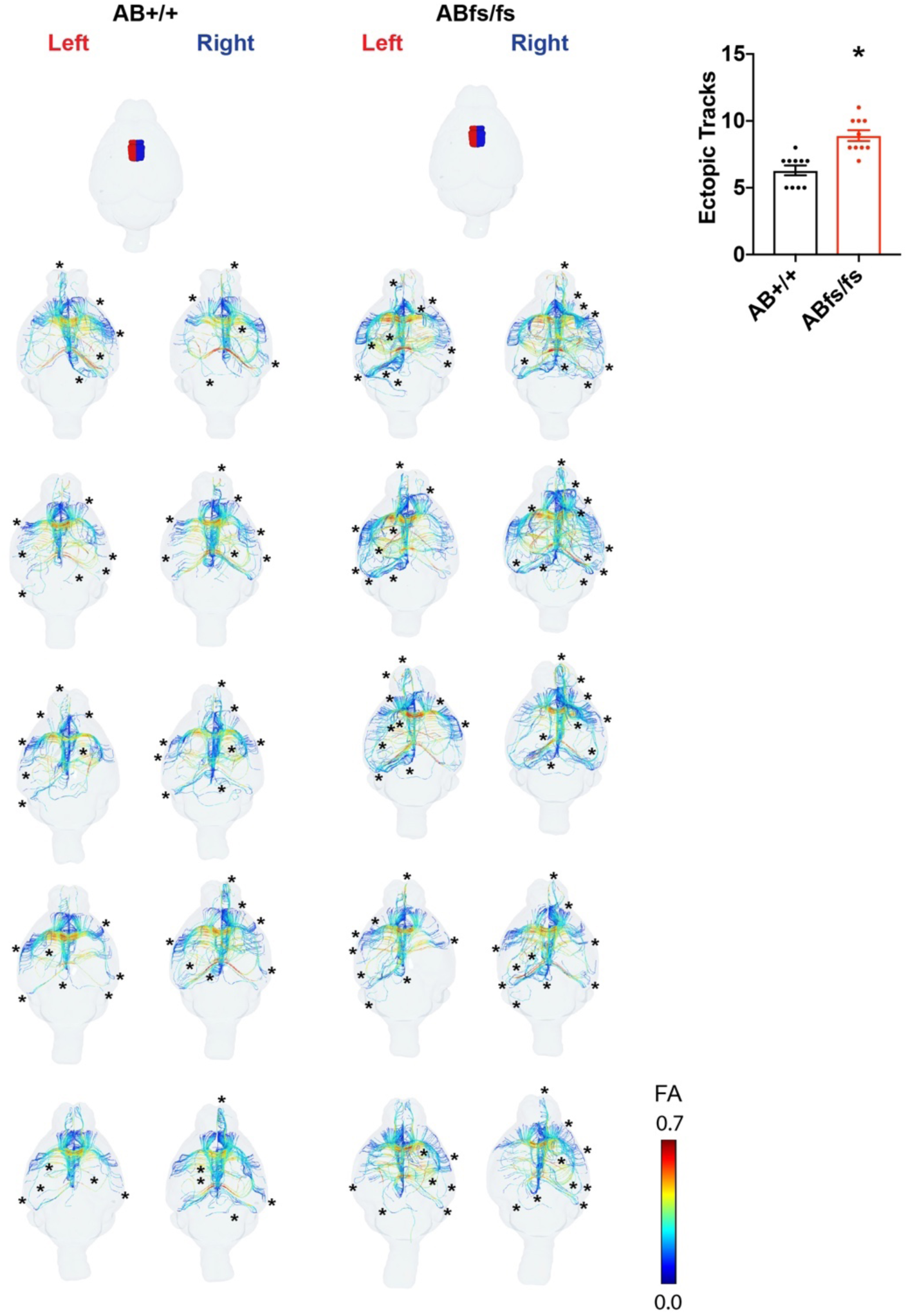
Tracts passing through the cingulate cortex area 24a of either the left or right hemisphere. Row 1: Rendering of cingulate cortex area 24a in left (red) or right (blue) hemisphere. Color based on FA values. * marks asymmetric tract. Quantification of ectopic (non-symmetrical) tracks. (n=5 mice/genotype). (Mean ±s.e.m.; t-test * P < 0.05).

**Extended Data Figure 3.**
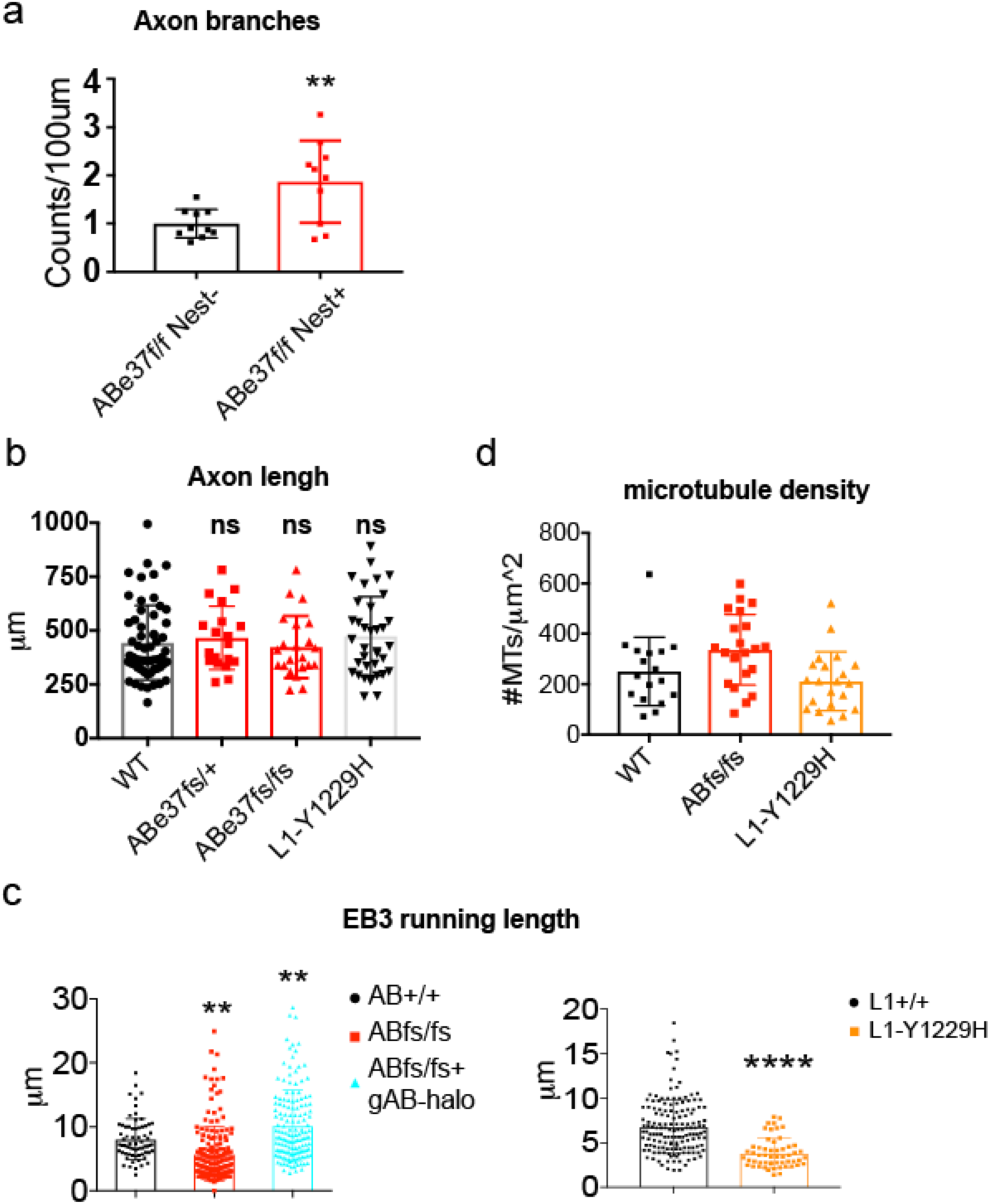
a. Axon branching quantification of 4-day old cultured hippocampal neurons (n=10). b. Axon length of cultured hippocampal neurons from indicated genotype (n=58, 19, 22, 35 from left to right in 3 cultures). c. Measurement of EB3 comet running length in live-imaged neurons (n=68,179,147, 151,55 from left to right in 3 cultures). d. Microtubule density in axons of the corpus callosum (n=17, 22, 21 axons in 2 mice/genotype). (Mean ±s.e.m.; a,c (right): t-test and b, c(left): one-way ANOVA followed by Dunnett’s multiple comparisons test; * P < 0.05; ** P < 0.01; *** P < 0.001; ****P<0.0001).

**Extended Data Figure 4.**
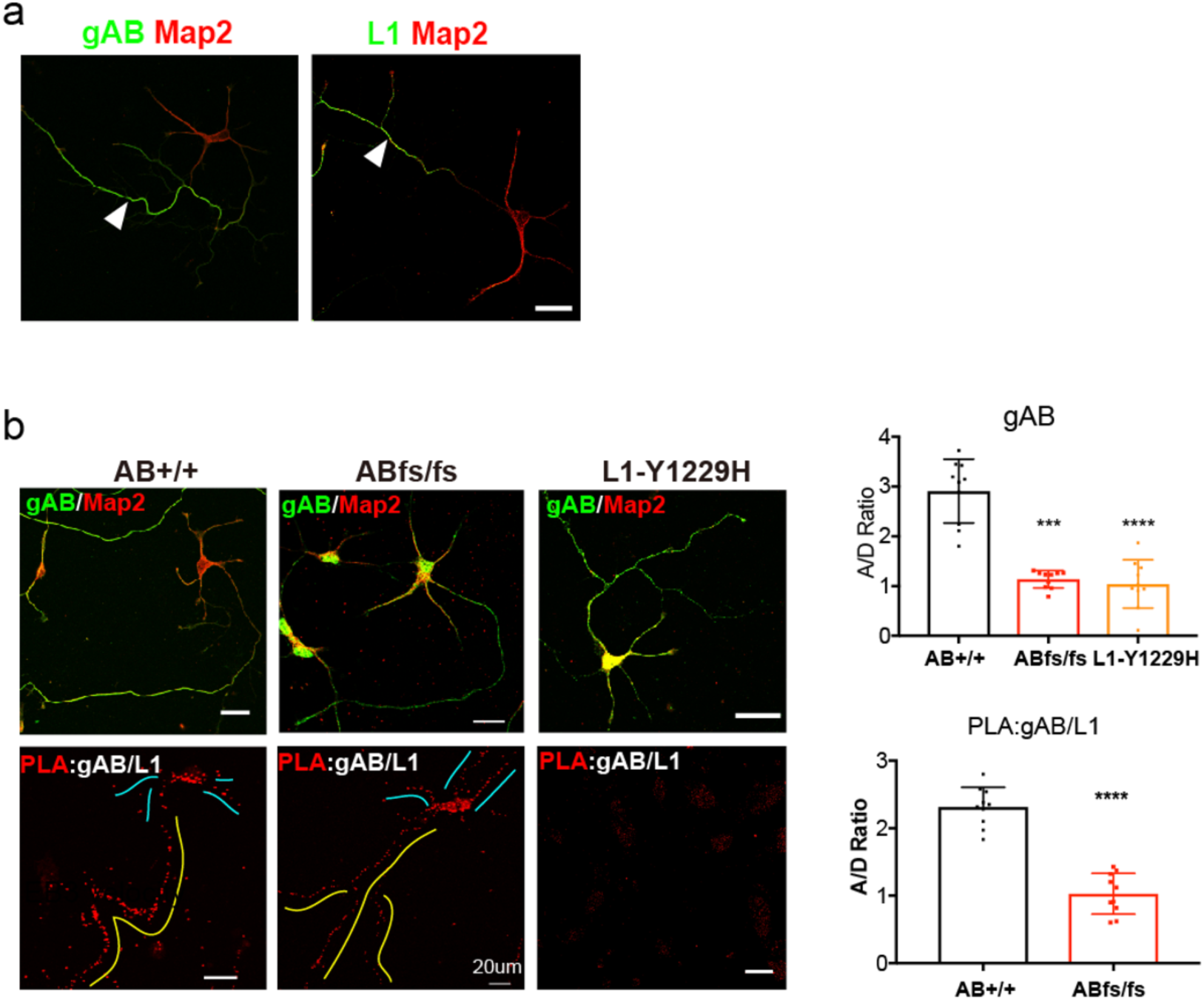
a. Immunostaining of gAB and L1CAM (green) in WT neurons. Map2 immunostaining (red) is used to identify dendrites. White arrow-head indicates axons. b. Immunostaining and proximity signal of giant ankB and L1CAM in cultured hippocampal neurons. Dendrites are highlighted with blue lines and the axon is highlighted with a yellow line. The axon to dendrite ratio of antibody staining and proximity signal are quantified. (n=10 cells/genotype in top panel n=10 cells/genotype in lower panel from left to right, scale bar: 20 µm). (Mean ±s.e.m; one-way ANOVA followed by Dunnett’s multiple comparisons test upper panel and t-test lower panel: ** P < 0.01; *** P < 0.001; **** P<0.0001).

**Extended Data Figure 5.**
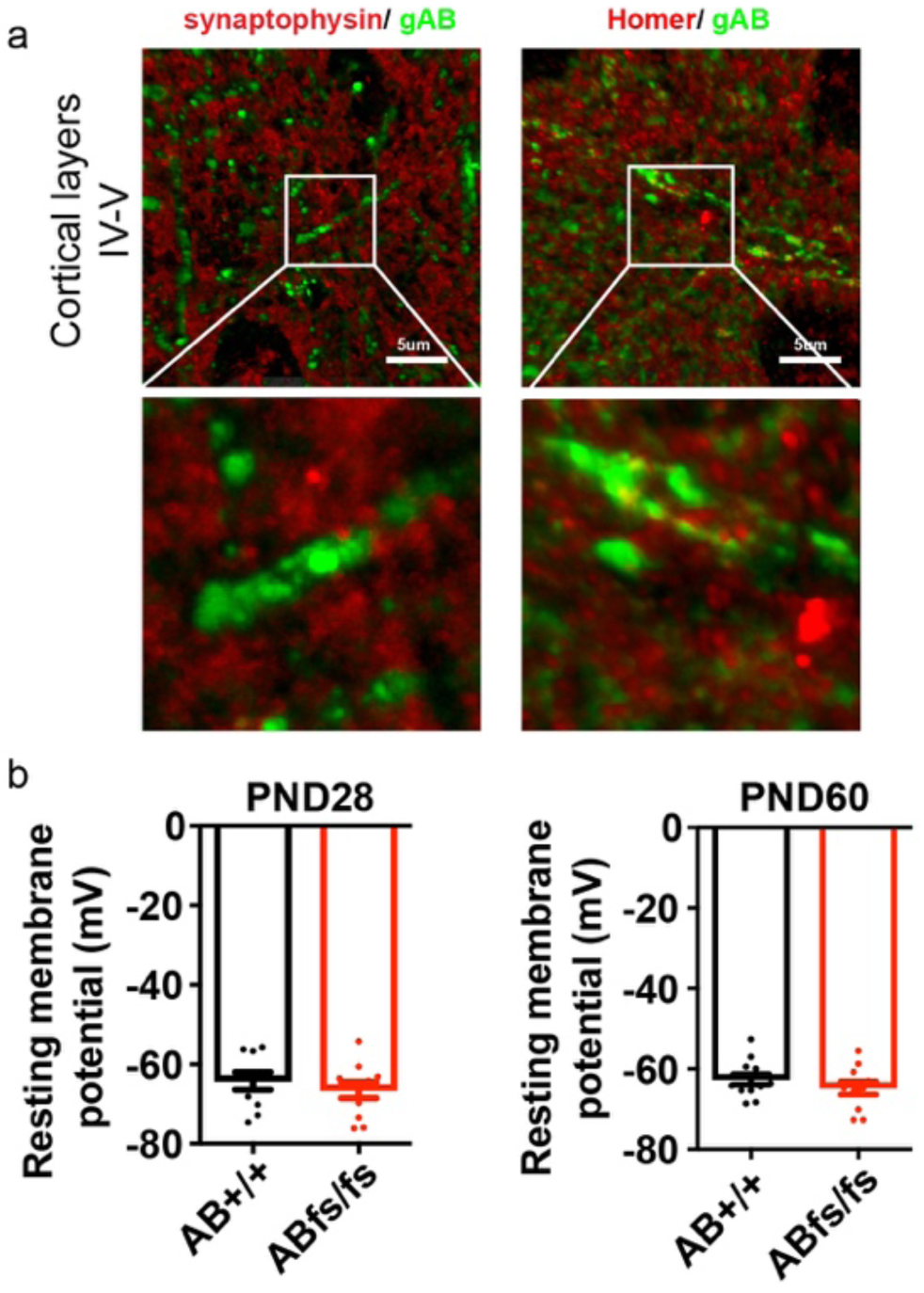
a. Immunostaining of gAB in green and synaptophysin or homer in red in the cortex (scale bar is 5µm). The region in the white box is enlarged and displayed below. b. Quantification of resting membrane potentials in neurons from PND28 and PND60 acute brain slices. N=8mice/genotype. (Mean ±s.e.m, no significant differences by t-test).

**Extended Data Figure 6a.**
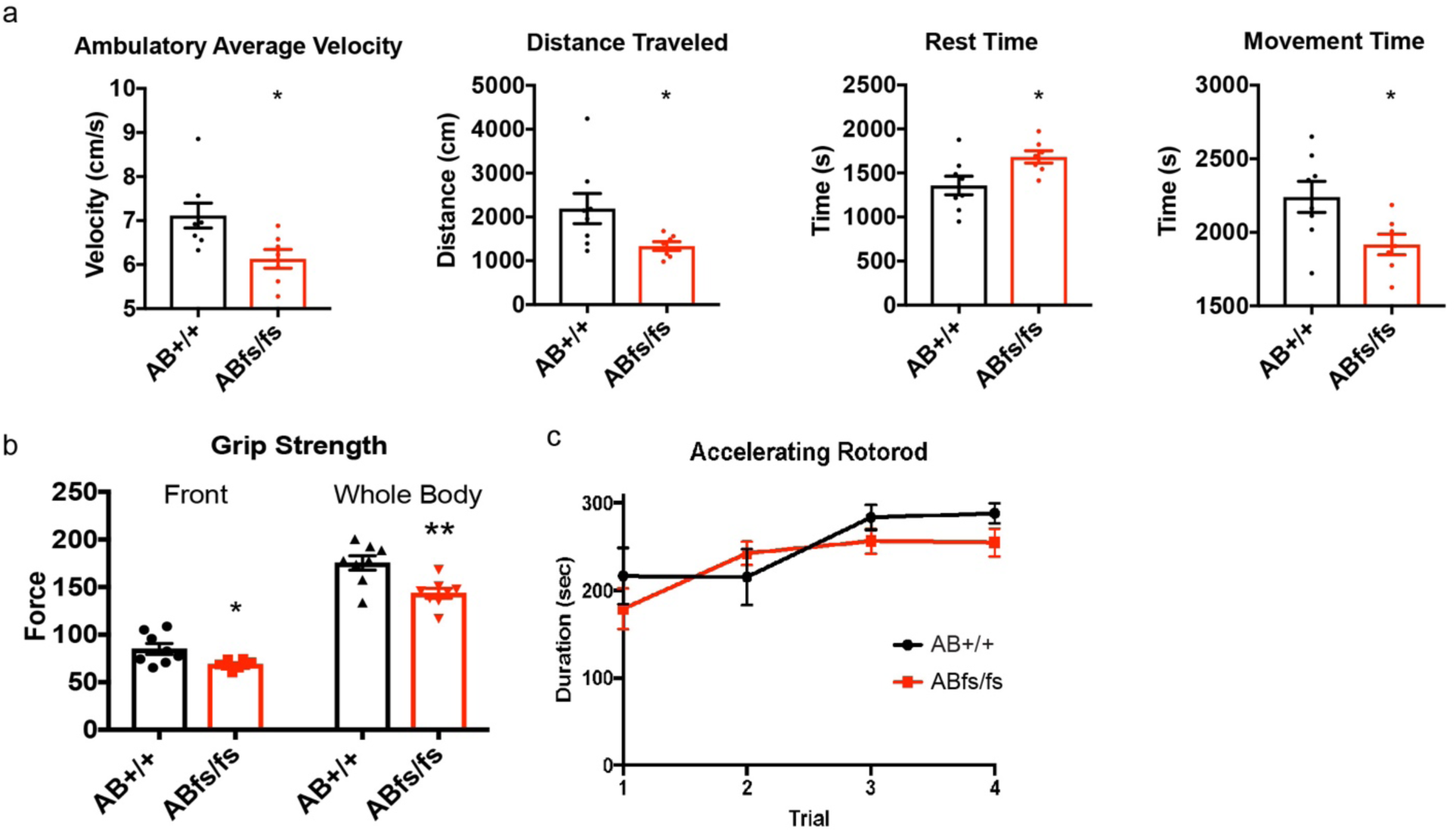
Motor Behavioral tests. a. Mean velocity, distance traveled, rest time, and movement time in the open field arena over 60 minutes. N=7-8mice/genotype; mean ±s.e.m. t-test *P<0.05. b. Grip strength of ABe37fs/fs mice. N=8mice/genotype; mean ±s.e.m. t-test *P<0.01, **P<0.001. c. Accelerating rotarod performance over consecutive trials. N=8mice/genotype; mean ±s.e.m. c, repeated measures ANOVA).

**Extended Data Figure 6b.**
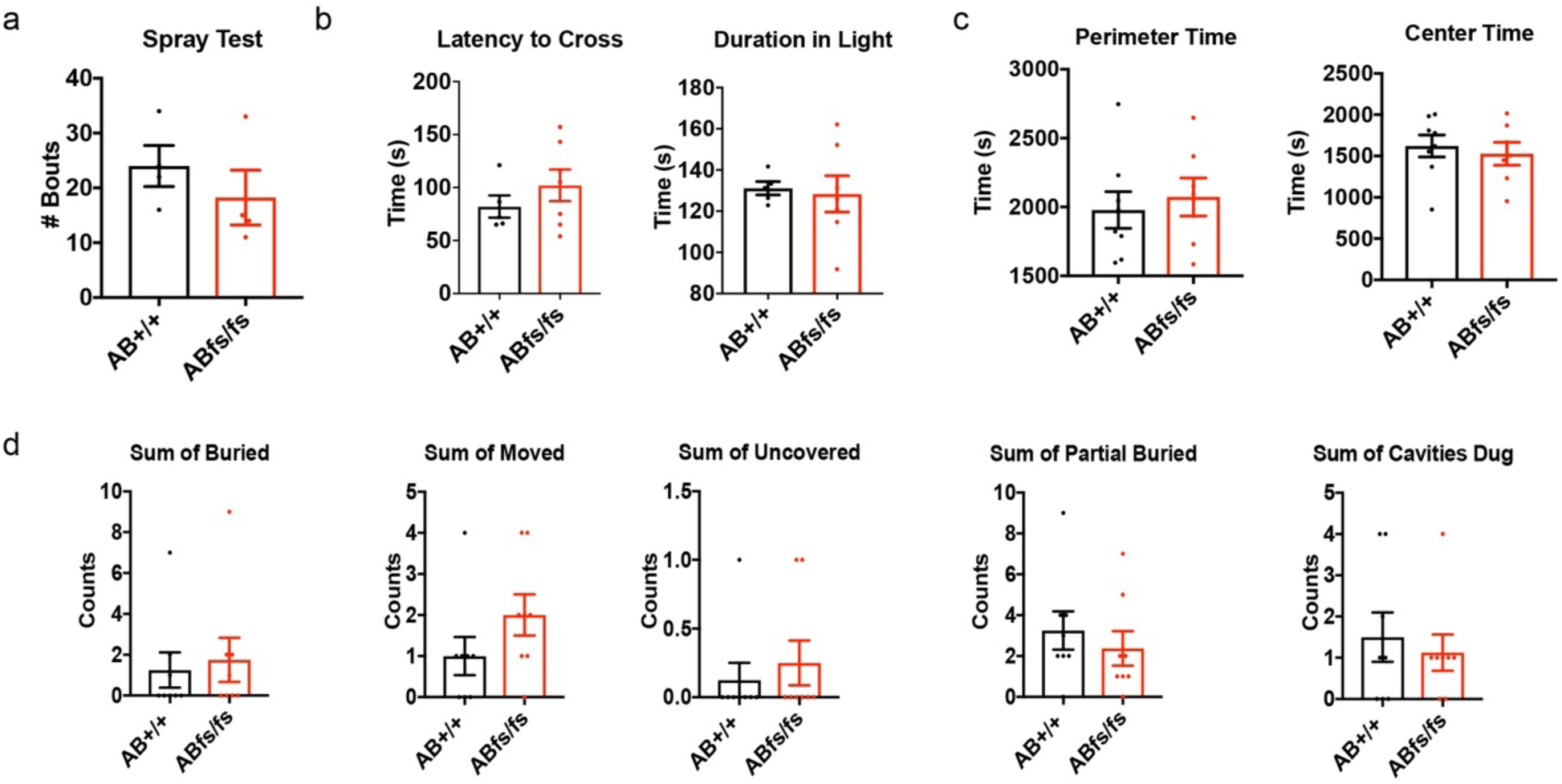
Repetitive behavior and anxiety-like responses. a. Number of grooming bouts in response to water spray. N=4mice/genotype. b. Latency to cross and time spent in the lighted chamber in the light-dark emergence test. N=5-8mice/genotype. c. Perimeter and center location time during a 1-hour open-field test. N=7-8 mice/genotype. d. Marble burying assay. N=8 mice/genotype. (Mean±s.e.m, t-test revealed no statistical difference for any test).

**Extended Data Figure 6c.**
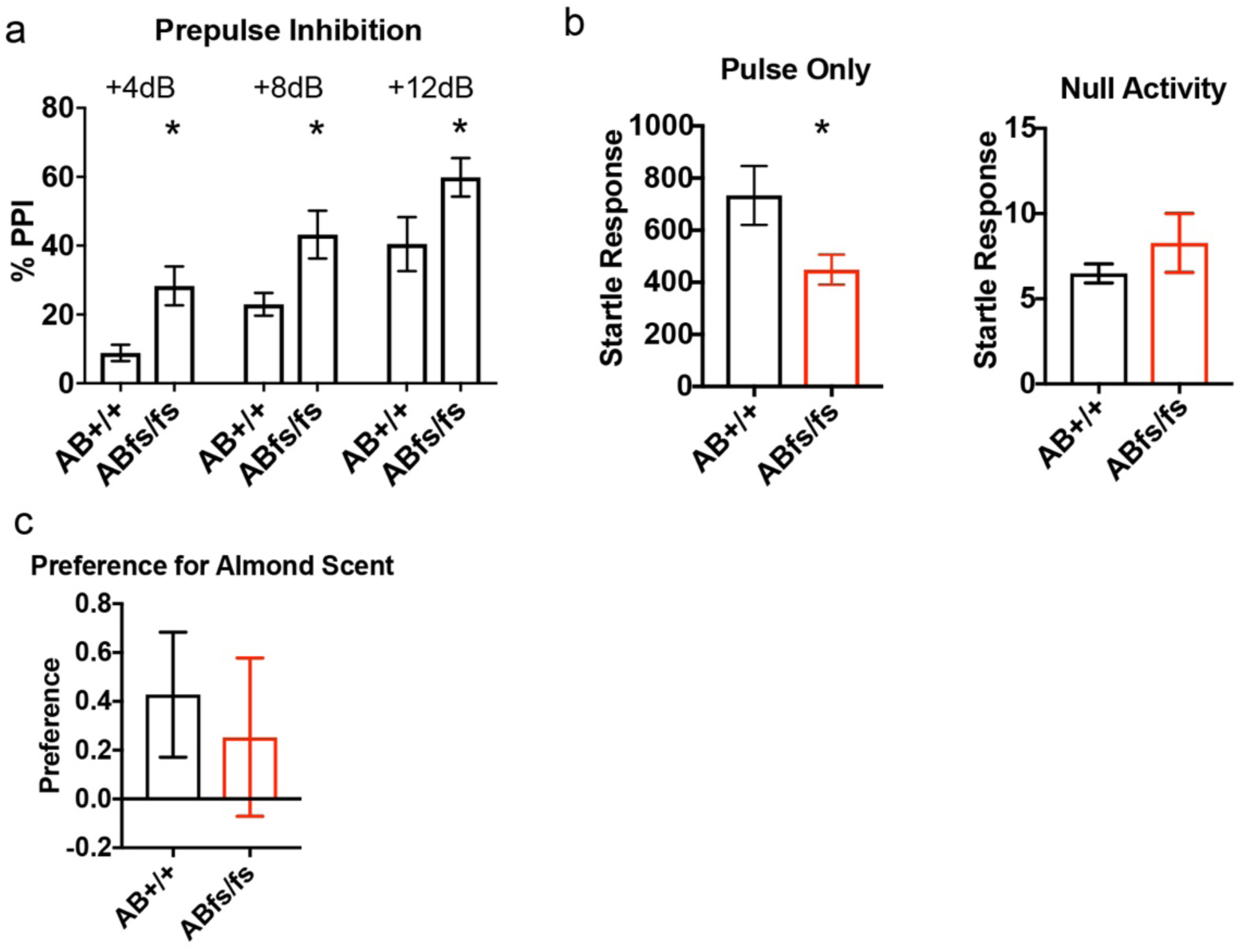
Sensorimotor gaiting and olfaction test. a. Auditory prepulse startle inhibition in ABe37fs/fs mice to the +4, 8, 12dB prepulses. b. Startle responses to the 120-dB stimulus and the baseline null activity. N=7-8mice/genotype. c. Olfaction test using almond extract. N=3mice/genotype. Mean ±s.e.m.; a, repeated measures ANOVA and b,c, t-test: * P >0.05.

**Extended Data Figure 6d.**
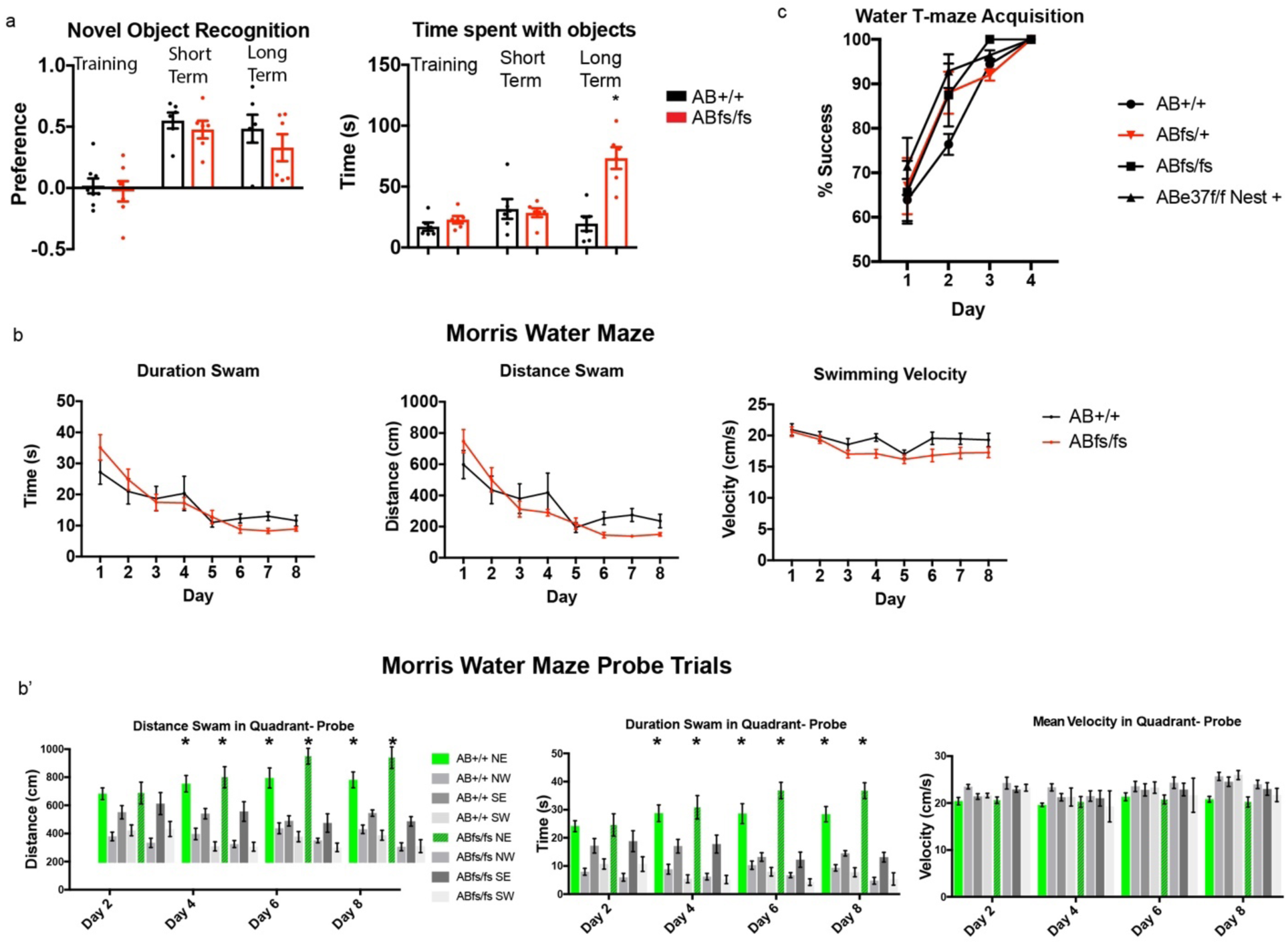
Cognitive assays in Ank2 mutant mice. a. Novel object recognition memory preference scores and time spent with objects in ABe37fs/fs mice displayed in training, and during tests of short- (30 min) and long-term (24 h) memory test. N=8mice/genotype. b. Acquisition performance in the Morris water maze task according to swim distance, swim time, and swim velocity in ABe37fs/fs mice averaged over 4 trials/day. N=7-8 mice/genotype. b’ Probe trials of acquisition performance in the Morris water maze task according to the same parameters in Abe37fs/fs mice. N=7-8mice/genotype. c. Acquisition phase in a water T-maze. Percent success calculated based on 4 trials/ day. N=7-9mice/genotype. (Mean ±s.e.m. repeated measures ANOVA).

**Extended Data Figure 7.**
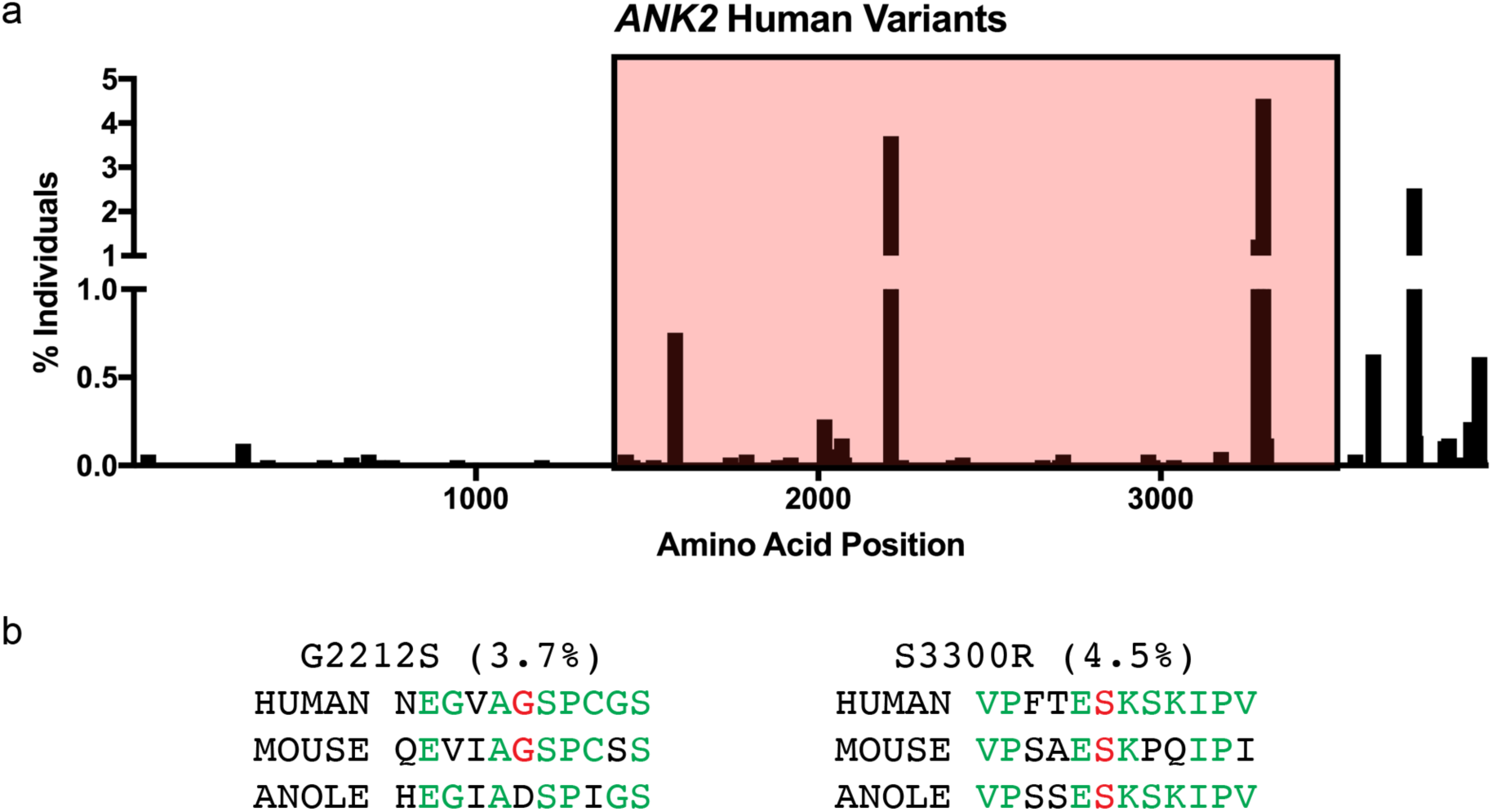
*ANK2* human variants with predicted deleterious polyphen scores in the exome variant server. a. Plot of % individuals with ANK2 mutation based on position of the mutation. Red shaded box indicates exon37 encoded region. Polyphen scores for missense mutations >0.7. b. Sequence comparison of missense sites for variants with %>3. % individuals affected in parenthesis. Red indicates mutated residue. Green indicates conserved residue.

**Extended Data Tables (attached separately as Excel files)**

Extended Data Table 1. Neurodevelopmental *ANK2* variants in humans

Extended Data Table 2. Gait analysis of ABe37fs/fs mice

Extended Data Table 3. MRI based brain volume analysis of ABe37fs/fs mice

Extended Data Table 4. Gene Expression analysis of ABe37fs DIV14 cultured neurons

Extended Data Table 5. GO analysis of gene expression from ABe37fs DIV14 cultured neurons

Extended Data Table 6. *ANK2* variants in humans from Exome Variant Server

## Acknowledgements

We thank E. Robinson and J. Hostettler for mouse husbandry and perfusion, S. Lalani for maintenance of primary neuronal culture, C. Guo (Janelia Gene Targeting and Transgenics Resource) for generating the ABe37fs mouse strain, Duke Transgenic Core for generating the ABe37floxed mouse strain, R. Rodriguez, F. Porka, and C. Means for behavioral experiment assistance (Duke Mouse Behavioral and Neuroendocrine Analysis Core), J. Christensen for coding and statistical analysis assistance, and J. Chen and Z. Bush for data analysis. We are grateful to the Duke Center for In Vivo Microscopy staff, in particular Y. Qi for perfusing the mice, G.A. Johnson, N. Delpratt, B.J. Anderson, and G. Coefer. We thank L. Cameron (Duke Light Microscopy Core Facility) for imaging advice for STED imaging and live-imaging. We thank D. Silver for suggesting the taxol rescue experiment. We thank S. Miller and R. Vancini of the Duke Electron Microscopy core for sample embedding and staining and V. Madden at the UNC Microscopy Service Laboratory for EM imaging. We thank R. Walder and B. Matsumoto for critical reading of the paper. VB was supported by R21MH115155, the Howard Hughes Medical Institute, and a George Barth Geller endowed professorship. KWC was supported by F31NS096848. HY was supported by MH112883. DTI imaging was funded by a Kahn Neurotechnology Development Grant to AB and VB and K01 AG041211 to AB. Duke In Vivo Microscopy core is supported in part by P41 EB015897.

## Author Contributions

The overall project was conceived by KWC and VB. Manuscript writing was done by KWC, VB, and RY. Genetic data were analyzed by SG, HS, CSA, LM, KWC, YHJ, and RY. DTI imaging was conceived by DL; AB acquired MRI and KWC, RY and AB analyzed data. The mechanistic cellular components were designed and interpreted by RY, KWC, and VB. DL performed initial proximity ligation experiments, which were confirmed and extended by RY and DW. DW and KWC performed immunoblotting and analyzed data. RY, KWC, DW performed synaptic counts. NK performed and analyzed electrophysiology experiments with HY. KWC performed behavior experiments and analysis with WCW. All authors contributed to data interpretation and critical revision of the manuscript.

## Contact for reagents and source data

Correspondence and requests for materials or additional source data should be directed to the corresponding authors, Vann Bennett or Kathryn Walder-Christensen.

